# Training deep neural density estimators to identify mechanistic models of neural dynamics

**DOI:** 10.1101/838383

**Authors:** Pedro J. Gonçalves, Jan-Matthis Lueckmann, Michael Deistler, Marcel Nonnenmacher, Kaan Öcal, Giacomo Bassetto, Chaitanya Chintaluri, William F. Podlaski, Sara A. Haddad, Tim P. Vogels, David S. Greenberg, Jakob H. Macke

**Affiliations:** Computational Neuroengineering, Department of Electrical and Computer Engineering, Technical University of Munich, Germany; Max Planck Research Group Neural Systems Analysis, Center of Advanced European Studies and Research (caesar), Bonn, Germany; Mathematical Institute, University of Bonn, Bonn, Germany; Centre for Neural Circuits and Behaviour, University of Oxford; Max Planck Institute for Brain Research, Frankfurt, Germany

## Abstract

Mechanistic modeling in neuroscience aims to explain observed phenomena in terms of underlying causes. However, determining which model parameters agree with complex and stochastic neural data presents a significant challenge. We address this challenge with a machine learning tool which uses deep neural density estimators— trained using model simulations— to carry out Bayesian inference and retrieve the full space of parameters compatible with raw data or selected data features. Our method is scalable in parameters and data features, and can rapidly analyze new data after initial training. We demonstrate the power and flexibility of our approach on receptive fields, ion channels, and Hodgkin–Huxley models. We also characterize the space of circuit configurations giving rise to rhythmic activity in the crustacean stomatogastric ganglion, and use these results to derive hypotheses for underlying compensation mechanisms. Our approach will help close the gap between data-driven and theory-driven models of neural dynamics.

## Introduction

New experimental technologies allow us to observe neurons, networks, brain regions and entire systems at unprecedented scale and resolution, but using these data to understand how behavior arises from neural processes remains a challenge. To test our understanding of a phenomenon, we often take to rebuilding it in the form of a computational model that incorporates the mechanisms we believe to be at play, based on scientific knowledge, intuition, and hypotheses about the components of a system and the laws governing their relationships. The goal of such mechanistic models is to investigate whether a proposed mechanism can explain experimental data, uncover details that may have been missed, inspire new experiments, and eventually provide insights into the inner workings of an observed neural or behavioral phenomenon [1–4]. Examples for such a symbiotic relationship between model and experiments range from the now classical work of Hodgkin and Huxley [5], to population models investigating rules of connectivity, plasticity and network dynamics [6–10], network models of inter-area interactions [11, 12], and models of decision making [13, 14].

A crucial step in building a model is adjusting its free parameters to be consistent with experimental observations. This is essential both for investigating whether the model agrees with reality and for gaining insight into processes which cannot be measured experimentally. For some models in neuroscience, it is possible to identify the relevant parameter regimes from careful mathematical analysis of the model equations. But as the complexity of both neural data and neural models increases, it becomes very difficult to find well-fitting parameters by inspection, and *automated* identification of data-consistent parameters is required.

Furthermore, to understand how a model quantitatively explains data, it is necessary to find not only the *best*, but *all* parameter settings consistent with experimental observations. This is especially important when modeling neural data, where highly variable observations can lead to broad ranges of data-consistent parameters. Moreover, many models in biology are inherently robust to some perturbations of parameters, but highly sensitive to others [3, 15], e.g. because of processes such as homeostastic regulation. For these systems, identifying the full range of data-consistent parameters can reveal how multiple distinct parameter settings give rise to the same model behavior [7, 16, 17]. Yet, despite the clear benefits of mechanistic models in providing scientific insight, identifying their parameters given data remains a challenging open problem that demands new algorithmic strategies.

The gold standard for automated parameter identification is *statistical inference*, which uses the likelihood *p*(**x**|***θ***) to quantify the match between parameters ***θ*** and data **x**. Likelihoods can be derived for purely statistical models commonly used in neuroscience [18–24], but are unavailable for most mechanistic models. Mechanistic models are designed to reflect knowledge about biological mechanisms, and not necessarily to be amenable to efficient inference: many mechanistic models are defined implicitly through stochastic computer simulations (e.g. a simulation of a network of spiking neurons), and likelihood calculation would require the ability to integrate over all potential paths through the simulator code. Similarly, a common goal of mechanistic modeling is to capture selected summary features of the data (e.g. a certain firing rate, bursting behavior, etc.‥), *not* the full dataset in all its details. The same feature (such as a particular average firing rate) can be produced by infinitely many realizations of the simulated process (such as a time-series of membrane potential). This makes it impractical to compute likelihoods, as one would have to average over all possible realizations which produce the same output.

Since the toolkit of statistical inference is inaccessible for mechanistic models, parameters are typically tuned ad-hoc (often through laborious, and subjective, trial-and-error), or by computationally expensive parameter search: a large set of models is generated, and grid search [25–27] or a genetic algorithm [28–31] is used to filter out simulations which do not match the data. However, these approaches require the user to define a heuristic rejection criterion on which simulations to keep (which can be challenging when observations have many dimensions or multiple units of measurement), and typically end up discarding most simulations. Furthermore, they lack the advantages of statistical inference, which provides principled approaches for handling variability, quantifying uncertainty, incorporating prior knowledge and integrating multiple data sources. Approximate Bayesian Computation (ABC) [32–34] is a parameter-search technique which aims to perform statistical inference, but still requires definition of a rejection criterion and struggles in high-dimensional problems. Thus, computational neuroscientists face a dilemma: either create carefully designed, highly interpretable mechanistic models (but rely on ad-hoc parameter tuning), or resort to purely statistical models offering sophisticated parameter inference but limited mechanistic insight.

Here we propose a new approach using machine learning to combine the advantages of mechanistic and statistical modeling. We present SNPE (Sequential Neural Posterior Estimation), a tool that rapidly identifies all mechanistic model parameters consistent with observed experimental data (or summary features). SNPE builds on recent advances in simulation-based Bayesian inference [35–38]: given observed experimental data (or summary features) **x**_*o*_, and a mechanistic model with parameters ***θ***, it expresses both prior knowledge and the range of data-compatible parameters through probability distributions. SNPE returns a posterior distribution *p*(***θ***|**x**_*o*_) which is high for parameters ***θ*** consistent with both the data **x**_*o*_ and prior knowledge, but approaches zero for ***θ*** inconsistent with either (Fig. 1).

**Figure 1.**
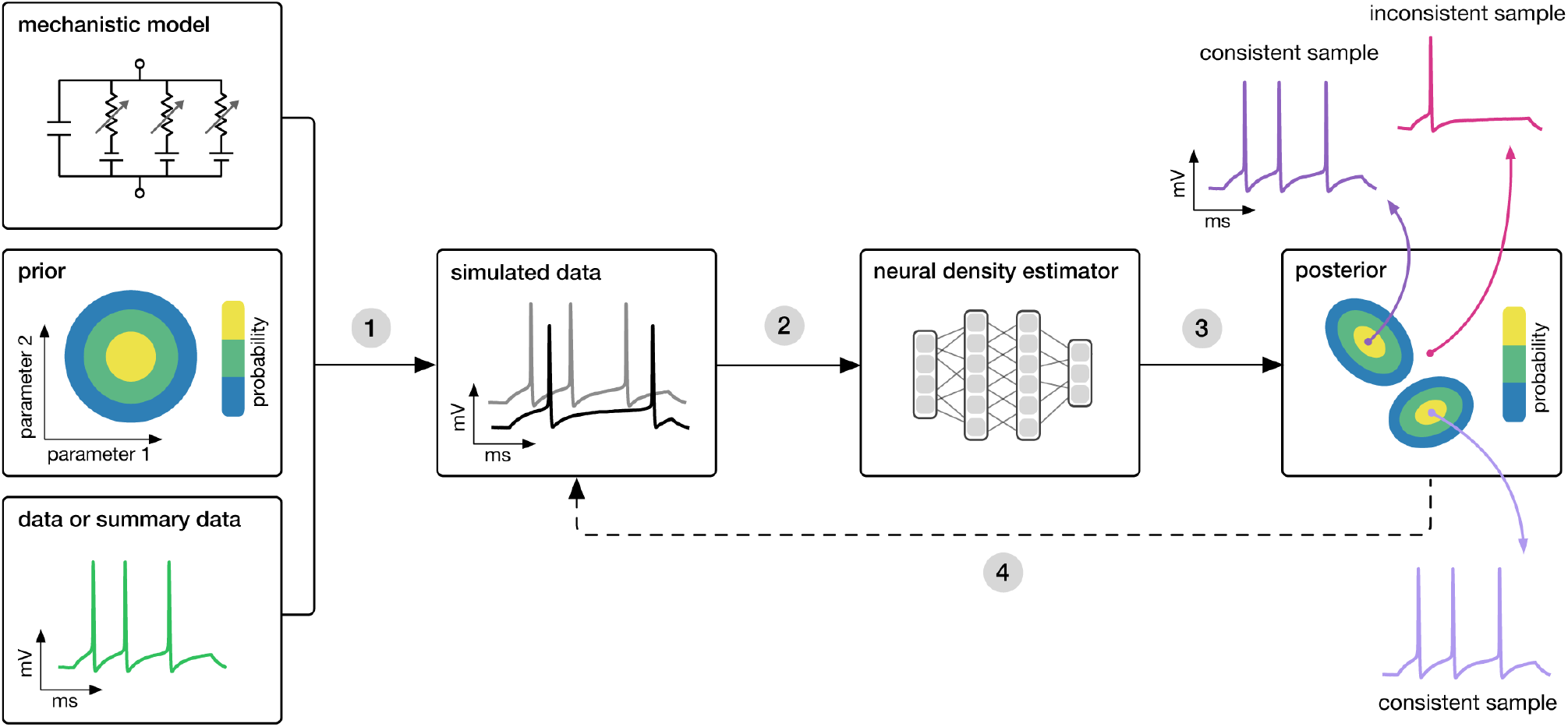
Goal: algorithmically identify mechanistic models which are consistent with data. Our algorithm (SNPE) takes three inputs: a candidate mechanistic model, prior knowledge or constraints on model parameters, and data (or summary statistics). SNPE proceeds by 1) sampling parameters from the prior and simulating synthetic datasets from these parameters, and 2) using a deep density estimation neural network to learn the (probabilistic) association between data (or data features) and underlying parameters, i.e. to learn statistical inference from simulated data. 3) This density estimation network is then applied to empirical data to derive the full space of parameters consistent with the data and the prior, i.e. the posterior distribution. High posterior probability is assigned to parameters which are consistent with both the data and the prior, low probability to inconsistent parameters. 4) If needed, an initial estimate of the posterior can be used to adaptively guide further simulations to produce data-consistent results.

Similar to parameter search methods, SNPE uses simulations instead of likelihood calculations, but instead of filtering out simulations, it uses *all* simulations to train a multi-layer artificial neural network to identify admissible parameters (Fig. 1). By incorporating modern deep neural networks for conditional density estimation [39, 40], it can capture the full *distribution* of parameters consistent with the data, even when this distribution has multiple peaks or lies on curved manifolds. Critically, SNPE decouples the design of the model and design of the inference approach, giving the investigator maximal flexibility to design and modify mechanistic models. Our method makes minimal assumptions about the model or its implementation, and can e.g. also be applied to non-differentiable models, such as networks of spiking neurons. Its only requirement is that one can run model simulations for different parameters, and collect the resulting synthetic data or summary features of interest.

While the theoretical foundations of SNPE were developed and tested using simple inference problems on small models [35–37], here we show that SNPE can scale to complex mechanistic models in neuroscience, provide an accessible and powerful implementation, and develop validation and visualization techniques for exploring the derived posteriors. We illustrate SNPE using mechanistic models expressing key neuroscientific concepts: beginning with a simple neural encoding problem with a known solution, we progress to more complex data types, large datasets and many-parameter models inaccessible to previous methods. We estimate visual receptive fields using many data features, demonstrate rapid inference of ion channel properties from high-throughput voltage-clamp protocols, and show how Hodgkin–Huxley models are more tightly constrained by increasing numbers of data features. Finally, we showcase the power of SNPE by using it to identify the parameters of a network model which can explain an experimentally observed pyloric rhythm in the stomatogastric ganglion [7]–in contrast to previous approaches, SNPE allows us to search over the full space of both single-neuron and synaptic parameters, allowing us to study the geometry of the parameter space, as well as to provide new hypotheses for which compensation mechanisms might be at play.

## Results

### Estimating stimulus-selectivity in linear-nonlinear encoding models

We first illustrate SNPE on linear-nonlinear (LN) encoding models, a special case of generalized linear models (GLMs). These are simple, commonly used phenomenological models for which likelihood-based parameter estimation is feasible [41–46], and which can be used to validate the accuracy of our approach, before applying SNPE to more complex models for which the likelihood is unavailable. We will show that SNPE returns the correct posterior distribution over parameters, that it can cope with high-dimensional observation data, that it can recover multiple solutions to parameter inference problems, and that it is substantially more simulation efficient than conventional rejection-based ABC methods.

An LN model describes how a neuron’s firing rate is modulated by a sensory stimulus through a linear filter ***θ***, often referred to as the *receptive field* [47, 48]. We first considered a model of a retinal ganglion cell (RGC) driven by full-field flicker (Fig. 2a). A statistic that is often used to characterize such a neuron is the *spike-triggered average* (STA) (Fig. 2a, right). We therefore used the STA, as well as the firing rate of the neuron, as input **x**_*o*_ to SNPE. (Note that, in the limit of infinite data, and for white noise stimuli, the STA will converge to the receptive field [42]–for finite, and non-white data, the two will in general be different.) Starting with random receptive fields ***θ***, we generated synthetic spike trains and calculated STAs from them (Fig. 2b). We then trained a neural conditional density estimator to recover the receptive fields from the STAs and firing rates (Fig. 2c). This allowed us to estimate the posterior distribution over receptive fields, i.e. to estimate which receptive fields are consistent with the data (and prior) (Fig. 2c). For LN models, likelihood-based inference is possible, allowing us to validate the SNPE posterior by comparing it to a reference posterior obtained via Markov Chain Monte Carlo (MCMC) sampling [45, 46]. We found that SNPE accurately estimates the posterior distribution (Supplementary Fig. 1 and Supplementary Fig. 2), and substantially outperforms Sequential Monte Carlo (SMC) ABC methods [34, 49] (Fig. 2d).

**Figure 2.**
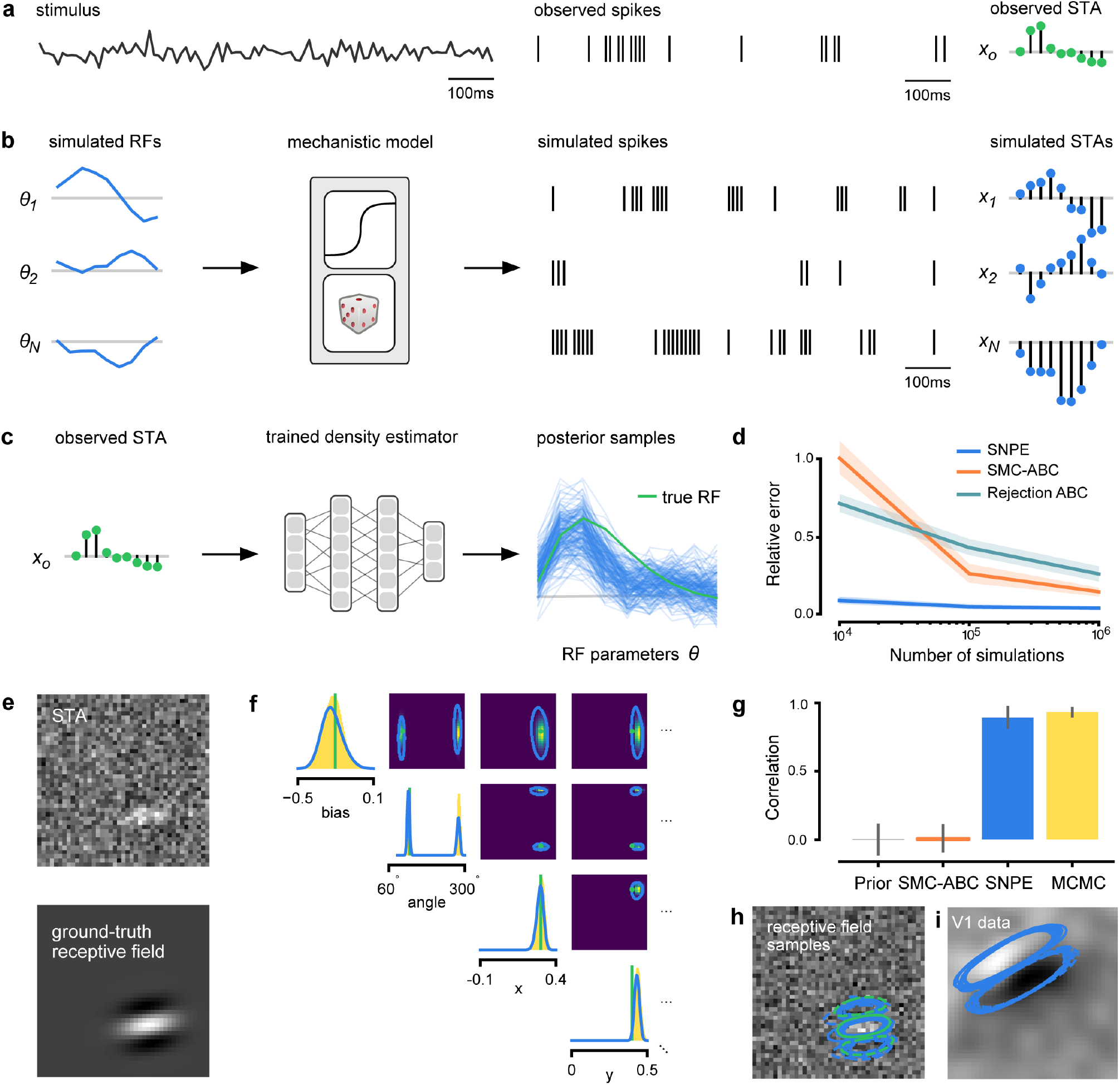
Estimating receptive fields in linear-nonlinear models of single neurons with statistical inference. (a) Schematic of a time-varying stimulus, associated observed spike train and resulting spike-triggered average (STA) (b) SNPE proceeds by first randomly generating simulated receptive fields *θ*, and using the mechanistic model (here an LN model) to generate simulated spike trains and simulated STAs. (c) These simulated STAs and receptive fields are then used to train a deep neural density estimator to identify the distribution of receptive fields consistent with a given observed STA **x**_*o*_. (d) Relative error in posterior estimation between SNPE and alternative methods (mean and 95%CI; 0 corresponds to perfect estimation, 1 to prior-level, details in Methods). (e) Example of spatial receptive field. We simulated responses and an STA of a LN-model with oriented receptive field. (f) We used SNPE to recover the distribution of receptive-field parameters. Univariate and pairwise marginals for four parameters of the spatial filter (MCMC, yellow histograms; SNPE, blue lines; full posterior in Supplementary Fig. 4). Non-identifiabilities of the Gabor parameterization lead to multimodal posteriors. (g) Average correlation (±SD) between ground-truth receptive field and receptive field samples from posteriors inferred with SMC-ABC, SNPE, and MCMC (which provides an upper bound given the inherent stochasticity of the data). (h) Posterior samples from SNPE posterior (SNPE, blue) compared to ground-truth receptive field (green; see panel (e)), overlaid on STA. (i) Posterior samples for V1 data; full posterior in Supplementary Fig. 5.

As a more challenging problem, we inferred the receptive field of a neuron in primary visual cortex (V1) [50, 51]. Using a model composed of a bias (related to the spontaneous firing rate) and a Gabor function with 8 parameters [52] describing the receptive field’s location, shape and strength, we simulated responses to 5-minute random noise movies of 41 × 41 pixels, such that the STA is high-dimensional, with a total of 1681 dimensions (Fig. 2e). This problem admits multiple solutions (as e.g. rotating the receptive field by 180°). As a result, the posterior distribution has multiple peaks (‘modes’). Starting from a simulation result **x**_*o*_ with known parameters, we used SNPE to estimate the posterior distribution *p*(***θ***|**x**_*o*_). To deal with the high-dimensional data **x**_*o*_ in this problem, we used a convolutional neural network (CNN), as this architecture excels at learning relevant features from image data [53, 54]. To deal with the multiple peaks in the posterior, we fed the CNN’s output into a mixture density network (MDN) [55], which can learn to assign probability distributions with multiple peaks as a function of its inputs (details in Methods). Using this strategy, SNPE was able to infer a posterior distribution that tightly enclosed the ground truth simulation parameters which generated the original simulated data **x**_*o*_, and matched a reference MCMC posterior (Fig. 2f, posterior over all parameters in Supplementary Fig. 4). For this challenging estimation problem with high-dimensional summary features, an SMC ABC algorithm with the same simulation-budget failed to identify the correct receptive fields (Fig. 2g) and posterior distributions (Supplementary Fig. 3). We also applied this approach to electrophysiological data from a V1 cell [51], identifying a sine-shaped Gabor receptive field consistent with the original spike-triggered average (Fig. 2i; posterior distribution in Supplementary Fig. 5).

### Functional diversity of ion channels: efficient high-throughput inference

We next show how SNPE can be efficiently applied to estimation problems in which we want to identify a large number of models for different observations in a database. We considered a flexible model of ion channels [57], which we here refer to as the *Omnimodel*. This model uses 8 parameters to describe how the dynamics of currents through non-inactivating potassium channels depend on membrane voltage (Fig. 3a). For various choices of its parameters ***θ***, it can capture 350 specific models in publications describing this channel type, cataloged in the IonChannelGenealogy (ICG) database [56]. We aimed to identify these ion channel parameters ***θ*** for each ICG model, based on 11 features of the model’s response to a sequence of 5 voltage clamp protocols, resulting in a total of 55 different characteristic features per model (Fig. 3b, see Methods for details).

**Figure 3.**
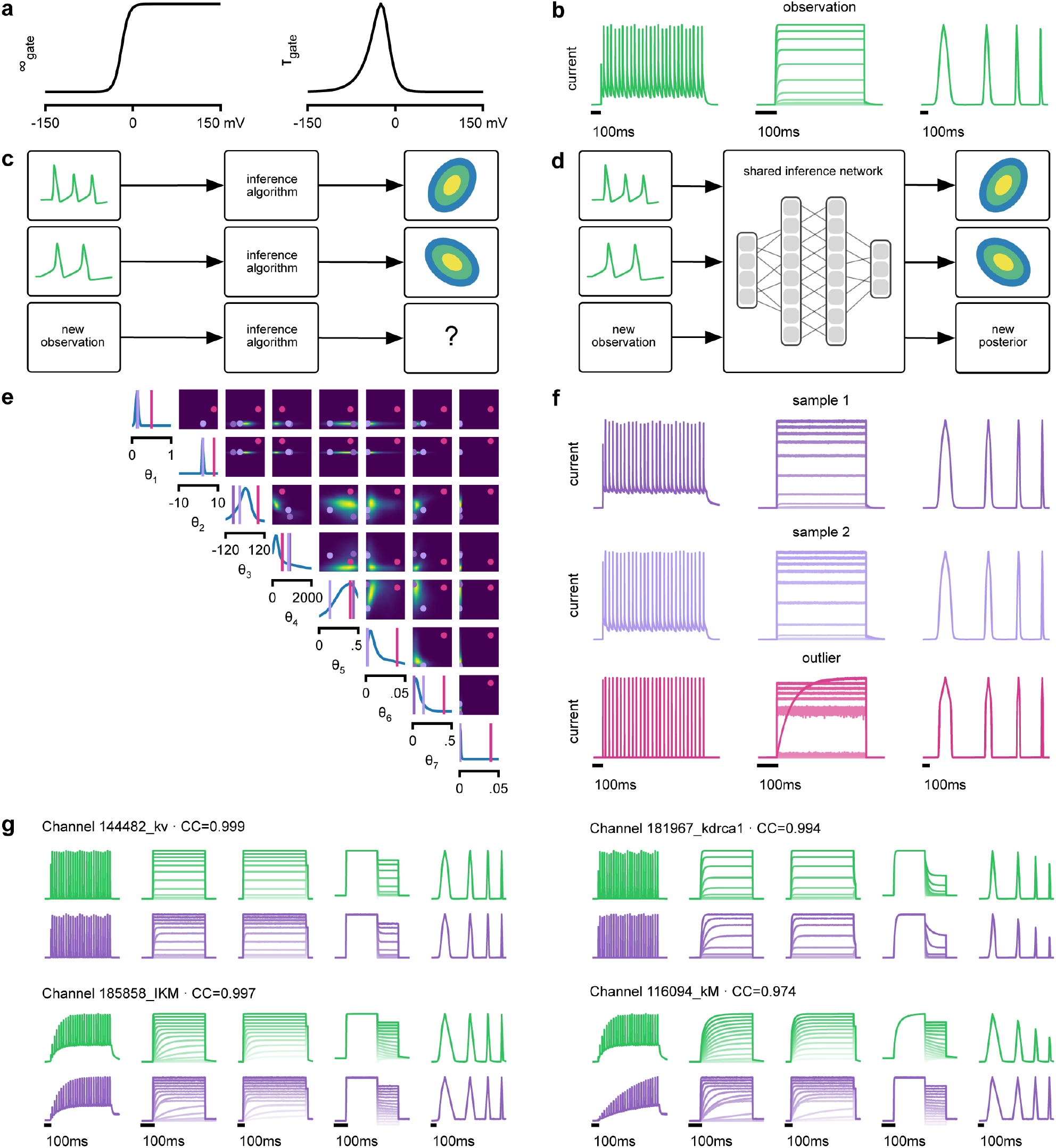
Inference on a database of ion-channel models. (a) We perform inference over the parameters of non-inactivating potassium channel models. Channel kinetics are described by steady-state activation curves, *∞*_gate_, and time-constant curves, *τ*_gate_. (b) Observation generated from a channel model from ICG database: normalized current responses to three (out of five) voltage-clamp protocols (action potentials, activation, and ramping). Details in [56]. (c) Classical approach to parameter identification: inference is optimized on each datum separately, requiring new computations for each new datum. (d) Amortized inference: an inference network is learned which can be applied to multiple data, enabling rapid inference on new data. (e) Posterior distribution over eight model parameters, *θ*_1_ to *θ*_8_. (f) Traces obtained by sampling from the posterior in (e). Purple: traces sampled from posterior, i.e. with high posterior probability. Magenta: trace from parameters with low probability. (g) Observations (green) and traces generated by posterior samples (purple) for four models from the database.

Because this model’s output is a typical format for functional characterization of ion channels both in simulations [56] and in high-throughput electrophysiological experiments [58–60], the ability to rapidly infer different parameters for many separate experiments is advantageous. Existing approaches for fitting deterministic models based on numerical optimization [57, 60] must repeat all computations anew for a new experiment or data point (Fig. 3c). However, for SNPE the only heavy computational tasks are carrying out simulations to generate training data, and training the neural network. We therefore reasoned that by training a network once using a large number of simulations, we could subsequently carry out rapid ‘amortized’ parameter inference on new data using a single pass through the network (Fig. 3d) [61, 62]. To test this idea, we used SNPE to train a neural network to infer the posterior from any data **x**. To generate training data, we carried out 1 million Omnimodel simulations, with parameters randomly chosen across ranges large enough to capture the models in the ICG database [56]. SNPE was run using a single round, i.e. it learned to perform inference for all data from the prior (rather than a specific observed datum). Generating these simulations took around 1000 CPU-hours and training the network 150 CPU-hours, but afterwards a full posterior distribution could be inferred for new data in less than 10 ms.

As a first test, SNPE was run on simulation data, generated by a previously published model of a non-inactivating potassium channel [63] (Fig. 3b). Simulations of the Omnimodel using parameter sets sampled from the obtained posterior distribution (Fig. 3e) closely resembled the input data on which the SNPE-based inference had been carried out, while simulations using ‘outlier’ parameter sets with low probability under the posterior generated current responses that were markedly different from the data **x**_*o*_ (Fig. 3f). Taking advantage of SNPE’s capability for rapid amortized inference, we further evaluated its performance on all 350 non-inactivating potassium channel models in ICG. In each case, we carried out a simulation to generate initial data from the original ICG model, used SNPE to calculate the posterior given the Omnimodel, and then generated a new simulation **x** using parameters sampled from the posterior (Fig. 3f). This resulted in high correlation between the original ICG model response and the Omnimodel response, in every case (>0.98 for more than 90% of models, see Supplementary Fig. 6). However, this approach was not able to capture all traces perfectly, as e.g. it failed to capture the shape of the onset of the bottom right model in Fig. 3g. Additional analysis of this example revealed that this example is not a failure of SNPE, but rather a limitation of the Omnimodel. Thus, SNPE can be used to reveal limitations of candidate models and aid the development of more verisimilar mechanistic models.

Calculating the posterior for all 350 ICG models took only a few seconds, and was fully automated, i.e. did not require user interactions. These results show how SNPE allows fast and accurate identification of biophysical model parameters on new data, and how SNPE can be deployed for applications requiring rapid automated inference, such as high-throughput screening-assays, closed-loop paradigms (e.g. for adaptive experimental manipulations or stimulus-selection), or interactive software tools.

### Hodgkin–Huxley model: stronger constraints from additional data features

The Hodgkin–Huxley (HH) model [5] of action potential generation through ion channel dynamics is a highly influential mechanistic model in neuroscience. A number of algorithms have been proposed for fitting HH models to electrophysiological data [25, 30, 31, 64–67], but [with the exception of 68] these approaches do not attempt to estimate the full posterior. Given the central importance of the HH model in neuroscience, we sought to test how SNPE would cope with this challenging non-linear model.

As previous approaches for HH models concentrated on reproducing specified features [e.g. the number of spikes, 65], we also sought to determine how various features provide different constraints. We considered the problem of inferring 8 biophysical parameters in a HH single-compartment model, describing voltage-dependent sodium and potassium conductances and other intrinsic membrane properties (Fig. 4a, left). We simulated the neuron’s voltage response to the injection of a square wave of depolarizing current, and defined the model output **x** used for inference as the number of evoked action potentials along with 6 additional features of the voltage response (Fig. 4a, right, details in Methods). We first applied SNPE to observed data **x**_*o*_ created by simulation from the model, calculating the posterior distribution using all 7 features in the observed data (Fig. 4b). The posterior contained the ground truth parameters in a high probability-region, as in previous applications, indicating the consistency of parameter identification. The variance of the posterior was narrower for some parameters than for others, indicating that the 7 data features constrain some parameters strongly (such as the potassium conductance), but others only weakly (such as the adaptation time constant). Additional simulations with parameters sampled from the posterior closely resembled the observed data **x**_*o*_, in terms of both the raw membrane voltage over time and the 7 data features (Fig. 4c, purple and green). Parameters with low posterior probability (outliers) generated simulations that markedly differed from **x**_*o*_ (Fig. 4c, magenta).

**Figure 4.**
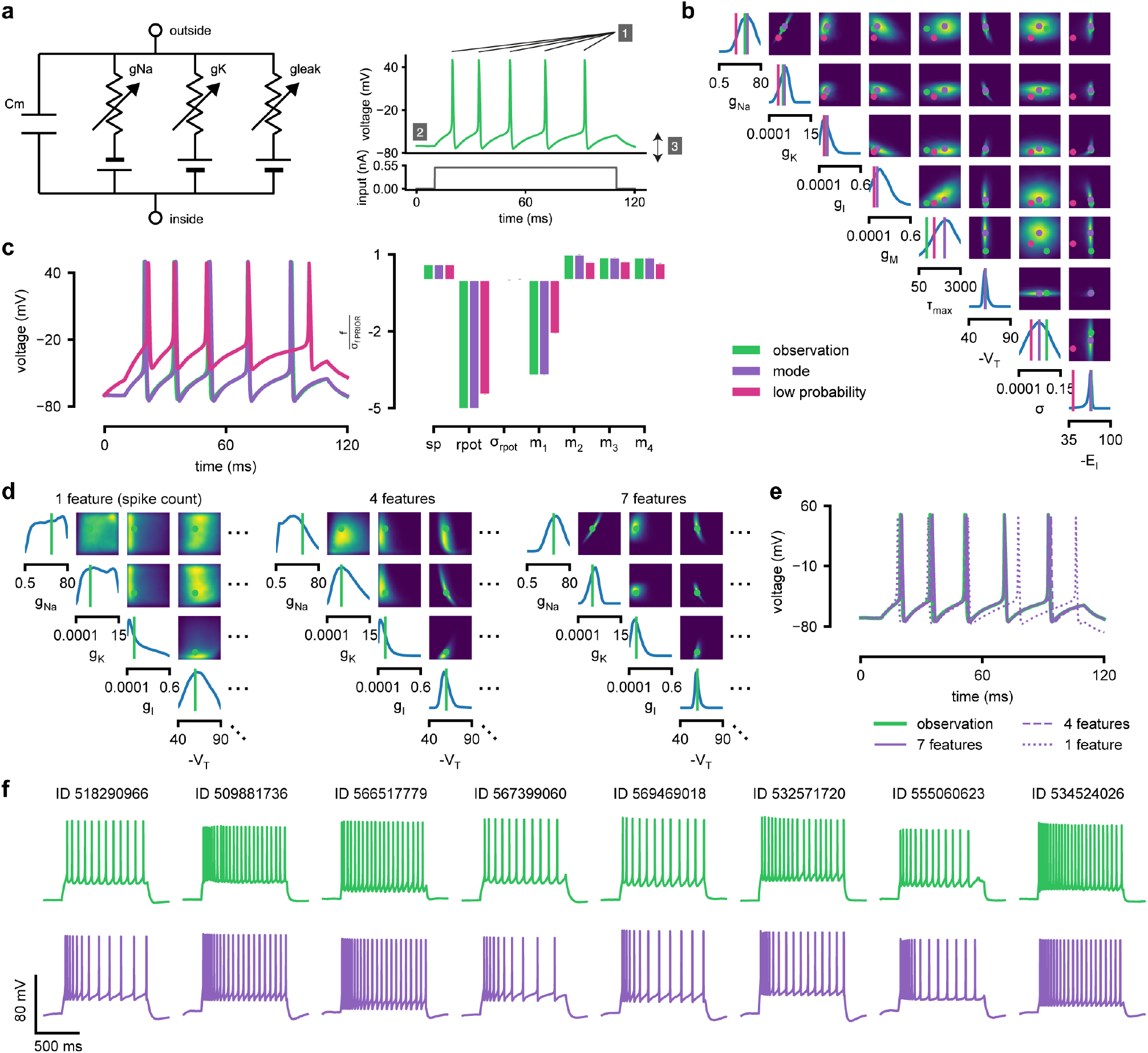
Inference for single compartment Hodgkin–Huxley model. (a) Circuit diagram describing the Hodgkin–Huxley model (left), and simulated voltage-trace given a current input (right). 3 out of 7 voltage features are depicted: (1) number of spikes, (2) mean resting potential and (3) standard deviation of the pre-stimulus resting potential. (b) Inferred posterior for 8 parameters given 7 voltage features. (c) Traces (left) and associated features *f* (right) for the desired output (observation), the mode of the inferred posterior, and a sample with low posterior probability. The voltage features are: number of spikes *sp*, mean resting potential *rpot*, standard deviation of the resting potential *σ*_rpot_, and the first 4 voltage moments, mean *m*_1_, standard deviation *m*_2_, skewness *m*_3_ and kurtosis *m*_4_. Each feature is normalized by *σ*_f PRIOR_, the standard deviation of the respective feature of simulations sampled from the prior. (d) Partial view of the inferred posteriors (4 out of 8 parameters) given 1, 4 and 7 features (full posteriors over 8 parameters in Supplementary Fig. 7). (e) Traces for posterior modes given 1, 4 and 7 features. Increasing the number of features leads to posterior traces that are closer to the observed data. (f) Observations from Allen Cell Types Database (green) and corresponding mode samples (purple). Posteriors in Supplementary Fig. 8.

Genetic algorithms are commonly used to fit parameters of deterministic biophysical models [28, 29, 31, 69]. While genetic algorithms can also return multiple data-compatible parameters, they do not perform inference (i.e. find the posterior distribution), and their outputs depend strongly on user-defined goodness-of-fit criteria. When comparing a state-of-the-art genetic algorithm [Indicator Based Evolutionary Algorithm, IBEA, 31, 70, 71] to SNPE, we found that the parameter-settings favoured by IBEA produced simulations whose summary features were as similar to the observed data as those obtained by SNPE high-probability samples (Supplementary Fig. 9). However, high-scoring IBEA parameters were concentrated in small regions of the posterior, i.e. IBEA did not identify the full space of data-compatible models.

To investigate how individual data features constrain parameters, we compared SNPE-estimated posteriors based 1) solely on the spike count, 2) on the spike count and 3 voltage-features, or 3) on all 7 features of **x**_*o*_. As more features were taken into account, the posterior became narrower and centered more closely on the ground truth parameters (Fig. 4d, Supplementary Fig. 7). Posterior simulations matched the observed data only in those features that had been used for inference (e.g. applying SNPE to spike counts alone identified parameters that generated the correct number of spikes, but for which spike timing and subthreshold voltage time course were off, Fig. 4e). For some parameters, such as the potassium conductance, providing more data features brought the peak of the posterior (the *posterior mode*) closer to the ground truth and also decreased uncertainty. For other parameters, such as *V_T_*, a parameter adjusting the spike threshold [65], the peak of the posterior was already close to the correct value with spike counts alone, but adding additional features reduced uncertainty. While SNPE can be used to study the effect of additional data features in reducing parameter uncertainty, this would not be the case for methods that only return a single best-guess estimate of parameters. These results show that SNPE can reveal how information from multiple data features imposes collective constraints on channel and membrane properties in the HH model.

We also inferred HH parameters for 8 *in vitro* recordings from the Allen Cell Types database using the same current-clamp stimulation protocol as in our model [72, 73] (Fig. 4f, Supplementary Fig. 8). In each case, simulations based on the SNPE-inferred posterior closely resembled the original data (Fig. 4f). We note that while inferred parameters differed across recordings, some parameters (the spike threshold, the density of sodium channels, the membrane reversal potential and the density of potassium channels) were consistently more strongly constrained than others (the intrinsic neural noise, the adaptation time constant, the density of slow voltage-dependent channels and the leak conductance) (Supplementary Fig. 8). Overall, these results suggest that the electrophysiological responses measured by this current-clamp protocol can be approximated by a single-compartment HH model, and that SNPE can identify the admissible parameters.

### Crustacean stomatogastric ganglion: sensitivity to perturbations

We next aimed to demonstrate how the full posterior distribution obtained with SNPE can lead to novel scientific insights. To do so, we used the pyloric network of the stomatogastric ganglion (STG) of the crab *Cancer borealis*, a well-characterized neural circuit producing rhythmic activity. In this circuit, similar network activity can arise from vastly different sets of membrane and synaptic conductances [7]. We first investigated whether data-consistent sets of membrane and synaptic conductances are connected in parameter space, as has been demonstrated for single neurons [75], and, second, which compensation mechanisms between parameters of this circuit allow the neural system to maintain its activity despite parameter variations. While this model has been studied extensively, answering these questions requires characterizing higher-dimensional parameter spaces than those accessed previously. We demonstrate how SNPE can be used to identify the posterior distribution over both membrane and synaptic conductances of the STG (31 parameters total) and how the full posterior distribution can be used to study the above questions at the circuit level.

For some biological systems, multiple parameter sets give rise to the same system behavior [7, 17, 76–79]. In particular, neural systems can be robust to specific perturbations of parameters [79–81], yet highly sensitive to others, properties referred to as *sloppiness* and *stiffness* [3, 15, 82, 83]. We studied how perturbations affect model output using a model [7] and data [74] of the pyloric rhythm in the crustacean stomatogastric ganglion (STG). This model describes a triphasic motor pattern generated by a well-characterized circuit (Fig. 5a). The circuit consists of two electrically coupled pacemaker neurons (anterior burster and pyloric dilator, AB/PD), modeled as a single neuron, as well as two types of follower neurons (lateral pyloric (LP) and pyloric (PY)), all connected through inhibitory synapses (details in Methods). Eight membrane conductances are included for each modeled neuron, along with 7 synaptic conductances, for a total of 31 parameters. This model has been used to demonstrate that virtually indistinguishable activity can arise from vastly different membrane and synaptic conductances in the STG [7, 17].

**Figure 5.**
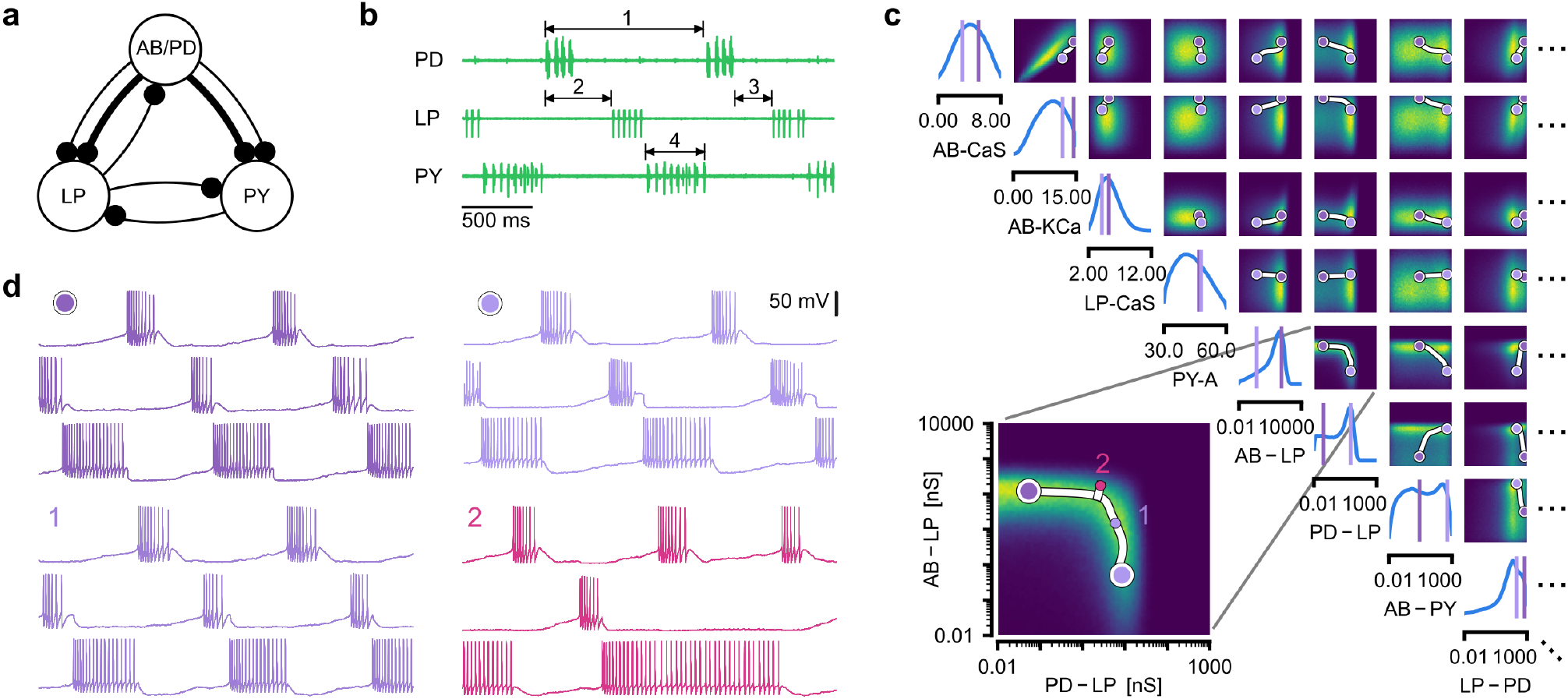
Identifying network models underlying an experimentally observed pyloric rhythm in the crustacean stomatogastric ganglion. (a) Simplified circuit diagram of the pyloric network from the stomatogastric ganglion. Thin connections are fast glutamatergic, thick connections are slow cholinergic. (b) Extracellular recordings from nerves of pyloric motor neurons of the crab *Cancer borealis* [74]. Numbers indicate some of the used summary features, namely cycle period (1), phase delays (2), phase gaps (3), and burst durations (4) (see Methods for details). (c) Posterior over 24 membrane and 7 synaptic conductances given the experimental observation shown in panel b (8 parameters shown, full posterior in Supplementary Fig. 10). Inset: magnified marginal posterior for the synaptic strengths AB to LP neuron vs. PD to LP neuron. (d) Identifying directions of sloppiness and stiffness. Two samples from the posterior both show similar network activity as the experimental observation (top left and top right), but have very different parameters (purple dots in panel c). Along the high-probability path between these samples, network activity is preserved (trace 1). When perturbing the parameters orthogonally off the path, network activity changes abruptly and becomes non-pyloric (trace 2).

We applied SNPE to an extracellular recording from the STG of the crab *Cancer borealis* [74] which exhibited pyloric activity (Fig. 5b), and inferred the posterior distribution over all 31 parameters based on 18 salient features of the voltage traces, including cycle period, phase delays, phase gaps, and burst durations (features in Fig. 5B, posterior in Fig. 5c, posterior over all parameters in Supplementary Fig. 10, details in Methods). Consistent with previous reports, the posterior distribution has high probability over extended value ranges for many membrane and synaptic conductances. To verify that parameter settings across these extended ranges are indeed capable of generating the experimentally observed network activity, we sampled two sets of membrane and synaptic conductances from the posterior distribution. These two samples have widely disparate parameters from each other (Fig. 5c, purple dots, details in Methods), but both exhibit activity highly similar to the experimental observation (Fig. 5d, top left and top right).

We then investigated the geometry of the parameter space producing these rhythms [16, 17]. First, we wanted to identify directions of sloppiness, and we were interested in whether parameter settings producing pyloric rhythms form a single connected region, as has been shown for single neurons [75], or whether they lie on separate ‘islands.’ Starting from the two above parameter settings showing similar activity, we examined whether they were connected by searching for a path through parameter space along which pyloric activity was maintained. To do this, we algorithmically identified a path lying only in regions of high posterior probability (Fig. 5c, white, details in Methods). Along the path, network output was tightly preserved, despite a substantial variation of the parameters (voltage trace 1 in Fig. 5d, Supplementary Fig. 11a,c). Second, we inspected directions of stiffness by perturbing parameters off the path. We applied perturbations that yield maximal drops in posterior probability (see Methods for details), and found that the network quickly produced non-pyloric activity (voltage trace 2, Fig. 5d) [82]. In identifying these paths and perturbations, we exploited the fact that SNPE provides a differentiable estimate of the posterior, as opposed to parameter search methods which provide only discrete samples.

Overall, these results show that the pyloric network can be robust to specific perturbations in parameter space, but sensitive to others, and that one can interpolate between disparate solutions while preserving network activity. This analysis demonstrates the flexibility of SNPE in capturing complex posterior distributions, and shows how the differentiable posterior can be used to study directions of sloppiness and stiffness.

### Predicting compensation mechanisms from posterior distributions

Experimental and computational studies have shown that stable neural activity can be maintained despite variable circuit parameters [7, 87, 88]. This behavior can emerge from two sources [87]: either, the variation of a certain parameter barely influences network activity at all, or alternatively, variations of several parameters influence network activity, but their effects compensate for one another. Here, we investigated these possibilities by using the posterior distribution over membrane and synaptic conductances of the STG.

We begin by drawing samples from the posterior and inspecting their pairwise histograms (i.e. the pairwise marginals, Fig. 6a, posterior over all parameters in Supplementary Fig. 10). Consistent with previously reported results [89], many parameters seem only weakly constrained and only weakly correlated (Fig. 6b). However, this observation does not imply that the parameters of the network do not have to be finely tuned: pairwise marginals are averages over many network configurations, where all other parameters may take on diverse values, which could disguise that each individual configuration is finely tuned. Indeed, when we sampled parameters independently from their posterior histograms, the resulting circuit configurations rarely produced pyloric activity, indicating that parameters have to be tuned relative to each other (Supplementary Fig. 12). This analysis also illustrates that the (common) approach of independently setting parameters can be problematic: although each parameter individually is in a realistic range, the network as a whole is not [90]. Finally, it shows the importance of identifying the full posterior distribution, which is far more informative than just finding individual parameters and assigning error bars.

**Figure 6.**
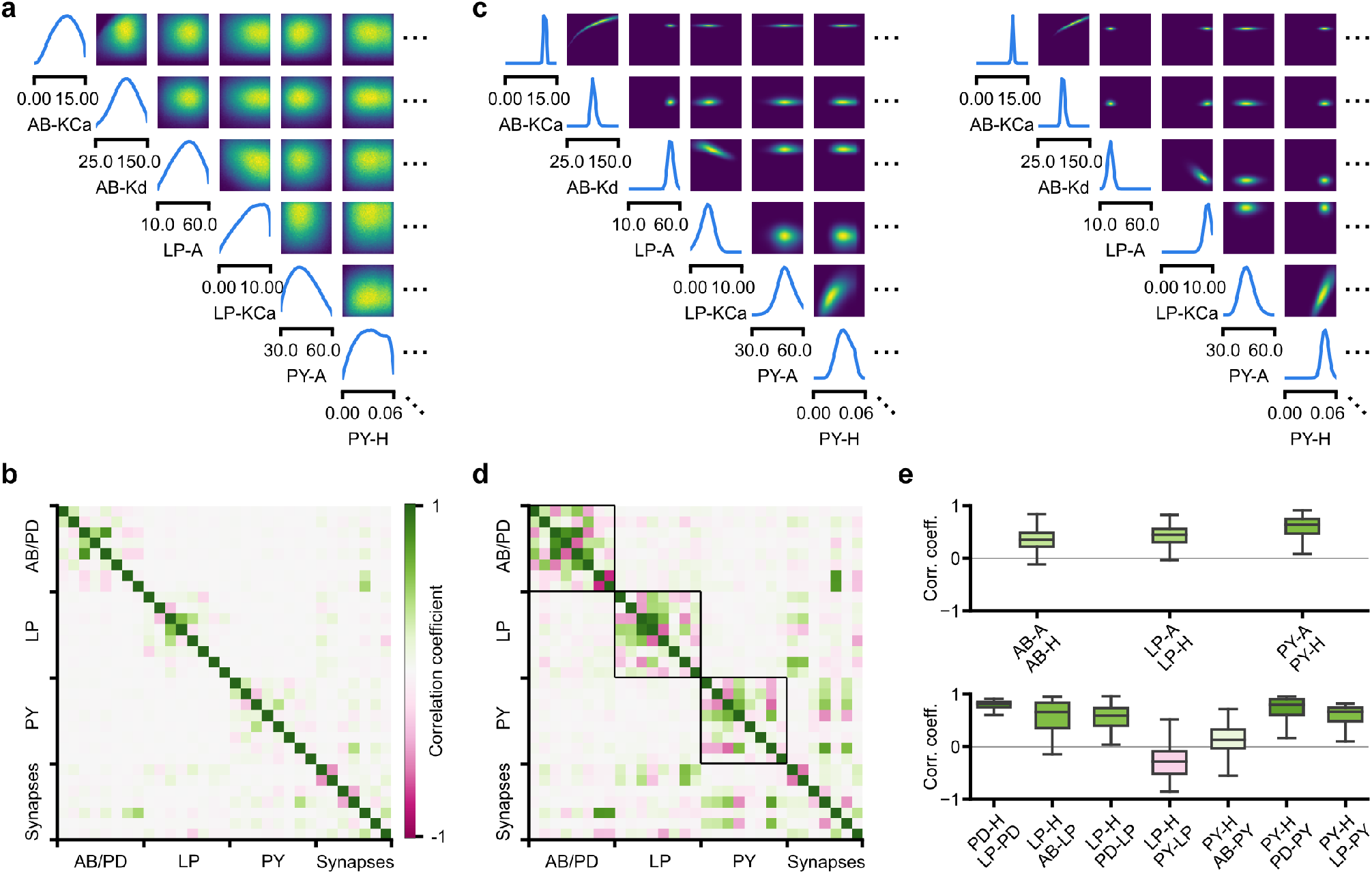
Predicting compensation mechanisms in the stomatogastric ganglion. (a) Inferred posterior. We show a subset of parameters which are weakly constrained (full posterior in Supplementary Fig. 10). Pyloric activity can emerge from a wide range of maximal membrane conductances, as the 1D and 2D posterior marginals cover almost the entire extent of the prior. (b) Correlation matrix, based on the samples shown in panel a. Almost all correlations are weak. Ordering of membrane and synaptic conductances as in Supplementary Fig. 10. (c) Conditional distributions given a particular circuit configuration: for the plots on the diagonal, we keep all but one parameter fixed. For plots above the diagonal, we keep all but two parameters fixed. The remaining parameter(s) are narrowly tuned, tuning across parameters is often highly correlated. When conditioning on a different parameter setting (right plot), the conditional posteriors change, but correlations are often maintained. (d) Conditional correlation matrix, averaged over 500 conditional distributions like the ones shown in panel c. Black squares highlight parameter-pairs within the same model neuron. (e) Consistency with experimental observations. Top: maximal conductance of the fast transient potassium current and the maximal conductance of the hyperpolarization current are positively correlated for all three neurons. This has also been experimentally observed in the PD and the LP neuron [84]. Bottom: the maximal conductance of the hyperpolarization current of the postsynaptic neuron can compensate the strength of the synaptic input, as experimentally observed in the PD and the LP neuron [85, 86]. The boxplots indicate the maximum, 75% quantile, median, 25% quantile, and minimum across 500 conditional correlations for different parameter pairs. Face color indicates mean correlation using the colorbar shown in panel b.

In order to investigate the need for tuning between pairs of parameters, we held all but two parameters constant at a given consistent circuit configuration (sampled from the posterior), and observed the network activity across different values of the remaining pair of parameters. We can do so by calculating the conditional posterior distribution (details in Methods), and do not have to generate additional simulations (as would be required by parameter search methods). Doing so has a simple interpretation: when all but two parameters are fixed, what values of the remaining two parameters can then lead to the desired network activity? We found that the desired pattern of pyloric activity can emerge only from narrowly tuned and often highly correlated combinations of the remaining two parameters, showing how these parameters can compensate for one another (Fig. 6c). When repeating this analysis across multiple network configurations, we found that these ‘conditional correlations’ are often preserved (Fig. 6c, left and right). This demonstrates that pairs of parameters can compensate for each other in a similar way, independently of the values taken by other parameters. This observation about compensation could be interpreted as an instance of modularity, a widespread underlying principle of biological robustness [91].

We calculated conditional correlations for each parameter pair using 500 different circuit configurations sampled from the posterior (Fig. 6d). Compared to correlations based on the pairwise marginals (Fig. 6b), these conditional correlations were substantially stronger. They were particularly strong across membrane conductances of the same neuron, but primarily weak across different neurons (black boxes in Fig. 6d).

Finally, we tested whether the conditional correlations were in line with experimental observations. For the PD and the LP neuron, it has been reported that overexpression of the fast transient potassium current (*I*_A_) leads to a compensating increase of the hyperpolarization current (*I*_H_), suggesting a positive correlation between these two currents [84, 92]. These results are qualitatively consistent with the positive conditional correlations between the maximal conductances of *I*_A_ and *I*_H_ for all three model neurons (Fig. 6e top). In addition, using the dynamic clamp, it has been shown that diverse combinations of the synaptic input strength and the maximal conductance of *I*_H_ lead to similar activity in the LP and the PD neuron [85, 86]. Consistent with these findings, the non-zero conditional correlations reveal that there can indeed be compensation mechanisms between the synaptic strength and the maximal conductance of *I*_H_ of the postsynaptic neuron (Fig. 6e bottom).

Overall, we showed how SNPE can be used to study parameter dependencies, and how the posterior distribution can be used to efficiently explore potential compensation mechanisms. We found that our method can predict compensation mechanisms which are qualitatively consistent with experimental studies. We emphasize that these findings would not have been possible with a direct grid-search over all parameters: defining a grid in a 31-dimensional parameter space would require more than 2^31^ >2 billion simulations, even if one were to use the coarsest-possible grid with only 2 values per dimension.

## Discussion

How can we build models which give insights into the causal mechanisms underlying neural or behavioral dynamics? The cycle of building mechanistic models, generating predictions, comparing them to empirical data, and rejecting or refining models has been of crucial importance in the empirical sciences. However, a key challenge has been the difficulty of identifying mechanistic models which can quantitatively capture observed phenomena. We suggest that a generally applicable tool to constrain mechanistic models by data would expedite progress in neuroscience. While many considerations should go into designing a model that is appropriate for a given question and level of description [2, 3, 93, 94], the question of whether and how one can perform statistical inference should not compromise model design. In our tool, SNPE, the process of model building and parameter inference are entirely decoupled. SNPE can be applied to *any* simulation-based model (requiring neither model nor summary features to be differentiable) and gives full flexibility on defining a prior. We illustrated the power of our approach on a diverse set of applications, highlighting the potential of SNPE to rapidly identify data-compatible mechanistic models, to investigate which data-features effectively constrain parameters, and to reveal shortcomings of candidate-models.

Finally, we used a model of the stomatogastric ganglion to show how SNPE can identify complex, high-dimensional parameter landscapes of neural systems. We analyzed the geometrical structure of the parameter landscape and confirmed that circuit configurations need to be finely tuned, even if individual parameters can take on a broad range of values. We showed that different configurations are connected in parameter space, and provided hypotheses for compensation mechanisms. These analyses were made possible by SNPE’s ability to estimate full parameter posteriors, rather than just constraints on individual parameters, as is common in many statistical parameter-identification approaches.

### Related work

SNPE builds on recent advances in machine learning, and in particular in density-estimation approaches to likelihood-free inference [35–37, 95, 96], reviewed in [38]. We here scaled these approaches to canonical mechanistic models of neural dynamics, and provided methods and software-tools for inference, visualization, and analysis of the resulting posteriors (e.g. the high-probability paths and conditional correlations presented here). The idea of learning inference networks on simulated data can be traced back to *regression-adjustment* methods in ABC [32, 97]. [35] first proposed to use expressive conditional density estimators in the form of deep neural networks [40, 55], and to optimize them sequentially over multiple rounds with cost-functions derived from Bayesian inference principles. Compared to commonly used rejection-based ABC methods [98, 99], such as MCMC-ABC [33], SMC-ABC [34, 100], Bayesian-Optimization ABC [101], or ensemble methods [102, 103], SNPE approaches do not require one to define a distance function in data space. In addition, by leveraging the ability of neural networks to learn informative features, they enable scaling to problems with high-dimensional observations, as are common in neuroscience and other fields in biology. We have illustrated this capability in the context of receptive field estimation, where a convolutional neural network extracts summary features from a 1681 dimensional spike-triggered average. Alternative likelihood-free approaches include *synthetic likelihood* methods [104–110], moment-based approximations of the posterior [111, 112], inference compilation [113, 114], and density-ratio estimation [115]. For some mechanistic models in neuroscience (e.g. for integrate-and-fire neurons), likelihoods can be computed via stochastic numerical approximations [66, 116, 117] or model-specific analytical approaches [64, 118–121].

Our approach is already finding its first applications in neuroscience–for example, [122] have used a variant of SNPE to constrain biophysical models of retinal neurons, with the goal of optimizing stimulation approaches for neuroprosthetics. Concurrently with our work, [123] developed an alternative approach to parameter identification for mechanistic models, and showed how it can be used to characterize neural population models which exhibit specific emergent computational properties. Both studies differ in their methodology and domain of applicability (see descriptions of underlying algorithms in our [36, 37] and their [124] prior work), as well in the focus of their neuroscientific contributions. Both approaches share the overall goal of using deep probabilistic inference tools to build more interpretable models of neural data. These complementary and concurrent advances will expedite the cycle of building, adjusting and selecting mechanistic models in neuroscience.

Finally, a complementary approach to mechanistic modeling is to pursue purely phenomenological models, which are designed to have favorable statistical and computational properties: these data-driven models can be efficiently fit to neural data [18–24, 41, 43] or to implement desired computations [125]. Although tremendously useful for a quantitative characterization of neural dynamics, these models typically have a large number of parameters, which rarely correspond to physically measurable or mechanistically interpretable quantities, and thus it can be challenging to derive mechanistic insights or causal hypotheses from them (but see e.g. [126–128]).

### Use of summary features

When fitting mechanistic models to data, it is common to target summary features to isolate specific behaviors, rather than the full data. For example, the spike shape is known to constrain sodium and potassium conductances [28, 29, 65]. When modeling population dynamics, it is often desirable to achieve realistic firing rates, rate-correlations and response nonlinearities [123, 129], or specified oscillations [7]. In models of decision making, one is often interested in reproducing psychometric functions or reaction-time distributions [130]. Choice of summary features might also be guided by known limitations of either the model or the measurement approach, or necessitated by the fact that published data are only available in summarized form. Several methods have been proposed to automatically construct informative summary features [131–133]. SNPE can be applied to, and might benefit from the use of summary features, but it also makes use of the ability of neural networks to automatically learn informative features in high-dimensional data. Thus, SNPE can also be applied directly to raw data (e.g. using recurrent neural networks [36]), or to high-dimensional summary features which are challenging for ABC approaches (Fig. 2). In all cases, care is needed when interpreting models fit to summary features, as choice of features can influence the results [131–133].

### Applicability and limitations

A key advantage of SNPE is its general applicability: it can be applied whenever one has a simulator that allows to stochastically generate model outputs from specific parameters. Furthermore, it can be applied in a fully ‘black-box manner’, i.e. does not require access to the internal workings of the simulator, its model equations, likelihoods or gradients. It does not impose any other limitations on the model or the summary features, and in particular does not require them to be differentiable. However, it also has limitations: first, current implementations of SNPE scale well to high-dimensional observations (~1000s dims, also see [37]), but scaling SNPE to even higher-dimensional parameter spaces (>30) is challenging (note that previous approaches were generally limited to dim < 10). Given that the difficulty of estimating full posteriors scales exponentially with dimensionality, this is an inherent challenge for all approaches that aim at full inference (in contrast to just identifying a single, or a few heuristically chosen parameter fits). Second, while it is a long-term goal for these approaches to be made fully automatic, our current implementation still requires choices by the user: as described in Methods, one needs to choose the type of the density estimation network, and specify settings related to network-optimisation, and the number of simulations and inference rounds. These settings depend on the complexity of the relation between summary features and model parameters, and the number of simulations that can be afforded. In the documentation accompanying our code-package, we provide examples and guidance. For small-scale problems, we have found SNPE to be robust to these settings. However, for challenging, high-dimensional applications, SNPE might currently require substantial user interaction. Third, the power of SNPE crucially rests on the ability of deep neural networks to perform density estimation. While deep nets have had ample empirical success, we still have an incomplete understanding of their limitations, in particular in cases where the mapping between data and parameters might not be smooth (e.g. near phase transitions). Fourth, when applying SNPE (or any other model-identification approach), validation of the results is of crucial importance, both to assess the accuracy of the inference procedure, as well as to identify possible limitations of the mechanistic model itself. In the example applications, we used several procedures for assessing the quality of the inferred posteriors. One common ingredient of these approaches is to sample from the inferred model, and search for systematic differences between observed and simulated data, e.g. to perform *posterior predictive checks* [36, 37, 100, 134, 135] (Fig. 2g, Fig. 3f,g, Fig. 4C, and Fig. 5d). There are challenges and opportunities ahead in further scaling and automating simulation-based inference approaches. However, in its current form, SNPE will be a powerful tool for quantitatively evaluating mechanistic hypotheses on neural data, and for designing better models of neural dynamics.

## Acknowledgments

We thank Mahmood S. Hoseini and Michael Stryker for sharing their data for Fig. 2, and Philipp Berens, Sean Bittner, Jan Boelts, John Cunningham, Richard Gao, Scott Linderman, Eve Marder, Iain Murray, George Papamakarios, Astrid Prinz, Auguste Schulz and Srinivas Turaga for discussions and/or comments on the manuscript. This work was supported by the German Research Foundation (DFG) through SFB 1233 ‘Robust Vision’, (276693517), SFB 1089 ‘Synaptic Microcircuits’ and SPP 2041 ‘Computational Connectomics’, the German Federal Ministry of Education and Research (BMBF, project ‘ADIMEM’, FKZ 01IS18052 A-D) to JHM, a Sir Henry Dale Fellowship by the Wellcome Trust and the Royal Society (WT100000; WFP and TPV), a Wellcome Trust Senior Research Fellowship (214316/Z/18/Z; TPV), a ERC Consolidator Grant (SYNAPSEEK; WPF and CC), and a UK Research and Innovation, Biotechnology and Biological Sciences Research Council (CC, UKRI-BBSRC BB/N019512/1.

## Methods

### Code availability

Code implementing SNPE is available at http://www.mackelab.org/delfi/.

### Simulation-based inference

To perform Bayesian parameter identification with SNPE, three types of input need to be specified:

1. A mechanistic model. The model only needs to be specified through a simulator, i.e. that one can generate a simulation result **x** for any parameters ***θ***. We do not assume access to the likelihood *p*(**x**|***θ***) or the equations or internals of the code defining the model, nor do we require the model to be differentiable. This is in contrast to many alternative approaches (including [123]), which require the model to be differentiable and to be implemented in a software code that is amenable to automatic differentiation packages. Finally, SNPE can both deal with inputs **x** which resemble ‘raw’ outputs of the model, or summary features calculated from data.
2. Observed data **x**_*o*_ of the same form as the results **x** produced by model simulations.
3. A prior distribution *p*(***θ***) describing the range of possible parameters. *p*(***θ***) could consist of upper and lower bounds for each parameter, or a more complex distribution incorporating mechanistic first principles or knowledge gained from previous inference procedures on other data.

For each problem, our goal was to estimate the posterior distribution *p*(***θ***|**x**_*o*_). To do this we used SNPE [35–37]. Setting up the inference procedure required three design choices:

1. A network architecture, including number of layers, units per layer, layer type (feedforward or convolutional), activation function and skip connections.
2. A parametric family of probability densities *q_ψ_*(***θ***) to represent inferred posteriors, to be used as conditional density estimator. We used either a mixture of Gaussians (MoG) or a masked autoregressive flow (MAF) [40]. In the former case, the number of components *K* must be specified; in the latter the number of *MADES* (Masked Autoencoder for Distribution Estimation) *n*_MADES_. Both choices are able to represent richly structured, and multimodal posterior distributions.
3. A simulation budget, i.e. number of rounds *R* and simulations per round *N_r_*.

We emphasize that SNPE is highly modular, i.e. that the the inputs (data, the prior over parameter, the mechanistic model), and algorithmic components (network architecture, probability density, optimization approach) can all be modified and chosen independently. This allows neuroscientists to work with models which are designed with mechanistic principles—and not convenience of inference—in mind. Furthermore, it allows SNPE to benefit from advances in more flexible density estimators, more powerful network architectures, or optimization strategies.

With the problem and inference settings specified, SNPE adjusts the network weights *ϕ* based on simulation results, so that *p*(***θ***|**x**) ≈ *q_F_* _(**x**,*ϕ*)_(***θ***) for any **x**. In the first round of SNPE simulation parameters are drawn from the prior *p*(***θ***). If a single round of inference is not sufficient, SNPE can be run in multiple rounds, in which samples are drawn from the version of 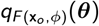 at the beginning of the round. After the last round, 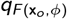 is returned as the inferred posterior on parameters ***θ*** given observed data **x**_*o*_. If SNPE is only run for a single round, then the generated samples only depend on the prior, but not on **x**_*o*_ : in this case, the inference network is applicable to any data (covered by the prior ranges), and can be used for rapid amortized inference.

SNPE learns the correct network weights *ϕ* by minimizing the objective function 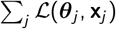 where the simulation with parameters ***θ***_*j*_ produced result **x**_*j*_. For the first round of SNPE 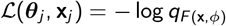, while in subsequent rounds a different loss function accounts for the fact that simulation parameters were not sampled from the prior. Different choices of the loss function for later rounds result in SNPE-A [35], SNPE-B [36] or SNPE-C algorithm [37]. To optimize the networks, we used ADAM with default settings [136].

The details of the algorithm are below:

**Algorithm 1:**
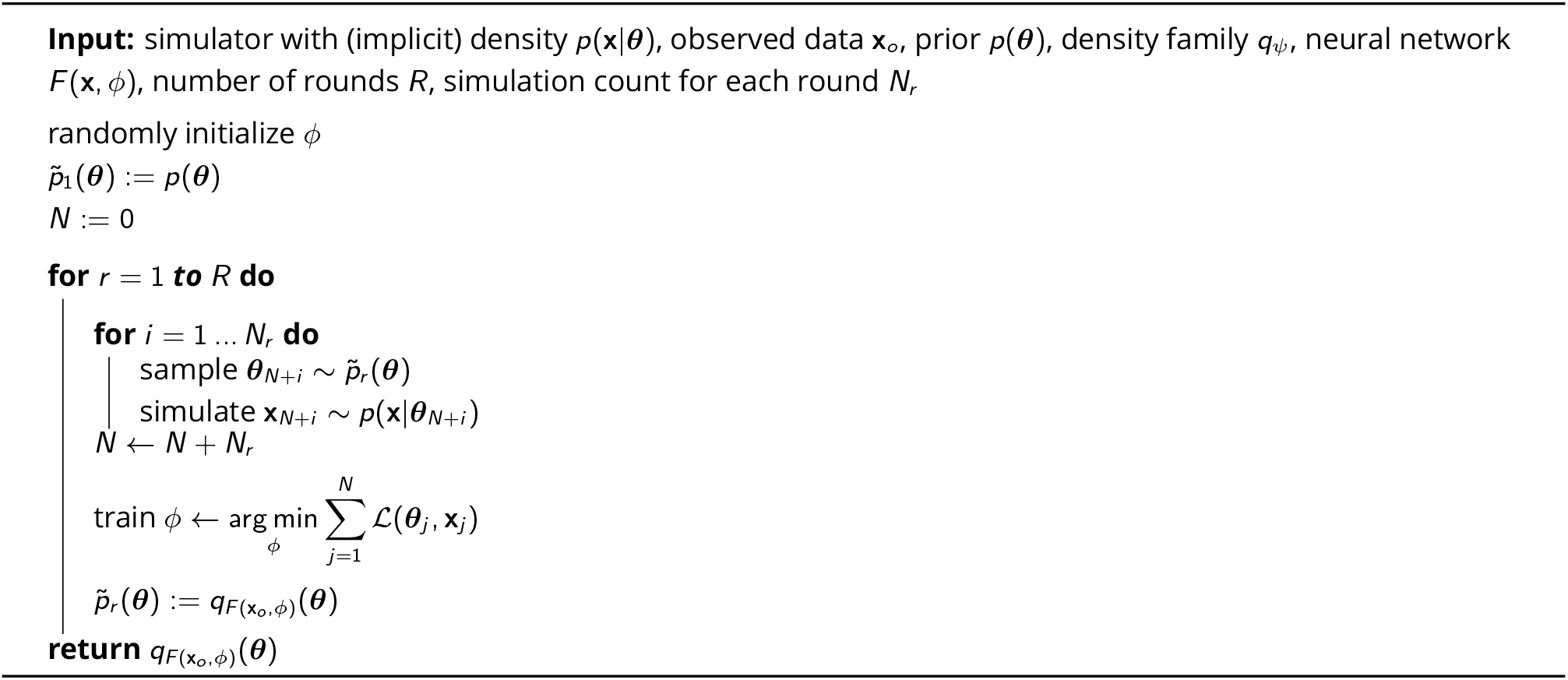
SNPE

### Linear-nonlinear encoding models

We used a Linear-Nonlinear (LN) encoding model (a special case of a generalized linear model, GLM, [18, 20, 41–44]) to simulate the activity of a neuron in response to a univariate time-varying stimulus. Neural activity *z_i_* was subdivided in *T* = 100 bins and, within each bin *i*, spikes were generated according to a Bernoulli observation model,

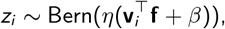

where **v**_*i*_ is a vector of white noise inputs between time bins *i* − 8 and *i*, **f** a length-9 linear filter, *β* is the bias, and *η*(·) = exp(·)/(1 + exp(·)) is the canonical inverse link function for a Bernoulli GLM. As summary features, we used the total number of spikes *N* and the spike-triggered average 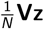, where **V** = [*v*_1_, *v*_2_, …, *v_T_*] is the so-called design matrix of size 9 × *T*. We note that the spike-triggered sum **Vz** constitutes sufficient statistics for this GLM, i.e. that selecting the STA and *N* together as summary features does not lead to loss of model relevant information over the full input-output dataset {**V**, **z**}. We used a Gaussian prior with zero mean and covariance matrix **Σ**_*β*_ = *σ*^2^(**F**^*T*^**F**)^*−*1^, where **F** encourages smoothness by penalizing the second-order differences in the vector of parameters [137].

For inference, we used a single round of 10000 simulations, and the posterior was approximated with a Gaussian distribution (***θ*** ∈ ℝ^10^, **x** ∈ ℝ^10^). We used a feedforward neural network with two hidden layers of 50 units each. We used a Polya Gamma Markov Chain Monte Carlo sampling scheme [45] to estimate a reference posterior.

In Fig. 2d, we compare the performance of SNPE with two classical ABC algorithms, rejection ABC and Sequential Monte Carlo ABC as a function of the number of simulations. We report the relative error in Kullback-Leibler divergence, which is defined as:

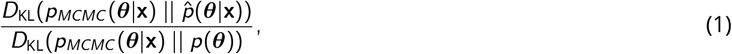

and which ranges between 0 (perfect recovery of the posterior) and 1 (estimated posterior no better than the prior). Here, *p_MCMC_* (***θ***|**x**) is the ground-truth posterior estimated via Markov Chain Monte Carlo sampling, 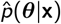 is the estimated posterior via SNPE, rejection ABC or Sequential Monte Carlo ABC, and *p*(***θ***) is the prior.

For the spatial receptive field model of a cell in primary visual cortex, we simulated the activity of a neuron depending on an image-valued stimulus. Neural activity was subdivided in bins of length Δ*t* = 0.025*s* and within each bin *i*, spikes were generated according to a Poisson observation model,

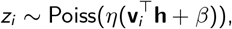

where **v**_*i*_ is the vectorized white noise stimulus at time bin *i*, **h** a 41 × 41 linear filter, *β* is the bias, and *η*(·) = exp(·) is the canonical inverse link function for a Poisson GLM. The receptive field **h** is constrained to be a Gabor filter:

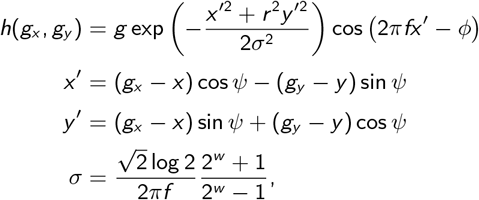

where (*g_x_*, *g_y_*) is a regular grid of 41 × 41 positions spanning the 2D image-valued stimulus. The parameters of the Gabor are gain *g*, spatial frequency *f*, aspect-ratio *r*, width *w*, phase *ϕ* (between 0 and *π*), angle *ψ* (between 0 and 2*π*) and location *x*, *y* (assumed within the stimulated area, scaled to be between −1 and 1). Bounded parameters were transformed with a log-, or logit-transform, to yield unconstrained parameters. After applying SNPE, we back-transformed both the parameters and the estimated posteriors in closed form, as shown in Fig. 2. We did not transform the bias *β*.

We used a factorizing Gaussian prior for the vector of transformed Gabor parameters

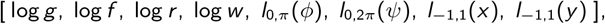

where transforms *l*_0,*π*_ (*X*) = log(*X/*(2*π* − *X*)), *l*_0,2*π*_ (*X*) = log(*X/*(*π* − *X*)), *l*_−1,1_(*X*) = log((*X* + 1)/(1 − *X*)) ensured the assumed ranges for the Gabor parameters *ϕ*, *ψ*, *x*, *y*. Our Gaussian prior had zero mean and standard deviations [0.5, 0.5, 0.5, 0.5, 1.9, 1.78, 1.78, 1.78]. We note that a Gaussian prior on a logit-transformed random variable logit*X* with zero mean and standard deviation around 1.78 is close to a uniform prior over the original variable *X*. For the bias *β*, we used a Gaussian prior with mean −0.57 and variance 1.63, which approximately corresponds to an exponential prior *exp*(*β*) ~ *Exp*(*λ*) with rate *λ* = 1 on the baseline firing rate exp(*β*) in absence of any stimulus.

The ground-truth parameters for the demonstration in Fig. 2 were chosen to give an asymptotic firing rate of 1Hz for 5 minutes stimulation, resulting in 299 spikes, and a signal-to-noise ratio of −12dB.

As summary features, we used the total number of spikes *N* and the spike-triggered average 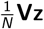, where **V** = [*v*_1_, *v*_2_, …, *v_T_*] is the stimulation video of length *T* = 300/Δ*t* = 12000. As for the GLM with a temporal filter, the spike-triggered sum **Vz** constitutes sufficient statistics for this GLM.

For inference, we applied SNPE-A with in total 2 rounds: an initial round serves to first roughly identify the relevant region of parameter space. Here we used a Gaussian distribution to approximate the posterior from 100000 simulations each. A second round then used a mixture of 8 Gaussian components to estimate the exact shape of the posterior from another 100000 simulations (***θ*** ∈ ℝ^9^, **x** ∈ ℝ^1682^). We used a convolutional network with 5 convolutional layers with 16 to 32 convolutional filters followed by two fully connected layers with 50 units each. The total number of spikes *N* within a simulated experiment was passed as an additional input directly to the fully-connected layers of the network. Similar to the previous GLM, this model has a tractable likelihood, so we use MCMC to obtain a reference posterior.

We applied this approach to extracelullar recordings from primary visual cortex of alert mice obtained using silicon microelectrodes in response to colored-noise visual stimulation. Experimental methods are described in [51].

### Comparison with Sequential Monte Carlo (SMC) ABC

In order to illustrate the competitive performance of SNPE, we obtained a posterior estimate with a classical ABC method, Sequential Monte Carlo (SMC) ABC [34, 49]. Likelihood-free inference methods from the ABC family require a distance function *d* (**x**_*o*_, **x**) between observed data **x**_*o*_ and possible simulation outputs **x** to characterize dissimilarity between simulations and data. A common choice is the (scaled) Euclidean distance *d* (**x**_*o*_, **x**) = ||**x** − **x**_*o*_ ||_2_. The Euclidean distance here was computed over 1681 summary features given by the spike-triggered average (one per pixel) and a single summary feature given by the ‘spike count’. To ensure that the distance measure was sensitive to differences in both STA and spike count, we scaled the summary feature ‘spike count’ to account for about 20% of the average total distance (other values did not yield better results). The other 80% were computed from the remaining 1681 summary features given by spike-triggered averages. To showcase how this situation is challenging for ABC approaches, we generated 10000 input-output pairs (***θ***_*i*_, **x**_*i*_) ~ *p*(**x**|***θ***)*p*(***θ***) with the prior and simulator used above, and illustrate the 10 STAs and spike counts with closest *d* (**x**_*o*_, **x**_*i*_) in Supplementary Fig. 3a. Spike counts were comparable to the observed data (299 spikes), but STAs are noise-dominated and the 10 ‘closest’ underlying receptive fields (orange contours) show substantial variability in location and shape of the receptive field. If even the ‘closest’ samples do not show any visible receptive field, then there is little hope that even an appropriately chosen acceptance threshold will yield a good approximation to the posterior. These findings were also reflected in the results from SMC-ABC with a total simulation budget of 10^6^ simulations (Fig. 3b). The estimated posterior marginals for ‘bias’ and ‘gain’ parameters show that the parameters related to the firing rate were constrained by the data **x**_*o*_, but marginals of parameters related to shape and location of the receptive field did not differ from the prior, highlighting that SMC-ABC was not able to identify the posterior distribution. The low correlations between the ground-truth receptive field and receptive fields sampled from SMC-ABC posterior further highlight the failure of SMC-ABC to infer the ground-truth posterior (Fig. 3c). Further comparisons of neural-density estimation approaches with ABC-methods can be found in the studies describing the underlying machine-learning methodologies [35, 37, 109].

### Ion channel models

We simulated non-inactivating potassium channel currents subject to voltage-clamp protocols as:

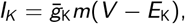

where *V* is the membrane potential, 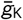 is the density of potassium channels, *E*_K_ is the reversal potential of potassium, and *m* is the gating variable for potassium channel activation. *m* is modeled according to the first-order kinetic equation

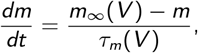

where *m*_∞_(*V*) is the steady-state activation, and *τ_m_* (*V*) the respective time constant. We used a general formulation of *m*_∞_(*V*) and *τ_m_* (*V*) [57], where the steady-state activation curve has 2 parameters (slope and offset) and the time constant curve has 6 parameters, amounting to a total of 8 parameters (*θ*_1_ to *θ*_8_):

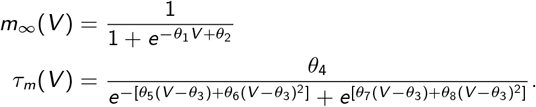

Since this model can be used to describe the dynamics of a wide variety of channel models, we refer to it as *Omnimodel*.

We modeled responses of the Omnimodel to a set of five voltage-clamp protocols described in [56]. Current responses were reduced to 55 summary features (11 per protocol). Summary features were coefficients to basis functions derived via Principal Components Analysis (PCA) (10 per protocol) plus a linear offset (1 per protocol) found via least-squares fitting. PCA basis functions were found by simulating responses of the non-inactivating potassium channel models to the five voltage-clamp protocols and reducing responses to each protocol to 10 dimensions (explaining 99.9% of the variance).

To amortize inference on the model, we specified a wide uniform prior over the parameters: 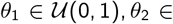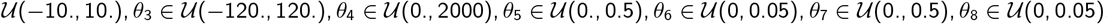.

For inference, we trained a shared inference network in a single round of 10^6^ simulations generated by sampling from the prior (***θ*** ∈ ℝ^8^, **x** ∈ ℝ^55^). The density estimator is a masked autoregressive flow (MAF) [40] with five MADES with [250,250] hidden units each.

We evaluated performance on 350 non-inactivating potassium ion channels selected from IonChannelGenealogy (ICG) by calculating the correlation coefficient between traces generated by the original model and traces from the Omnimodel using the posterior mode.

### Single-compartment Hodgkin–Huxley neurons

We simulated a single-compartment Hodgkin–Huxley type neuron with channel kinetics as in [65],

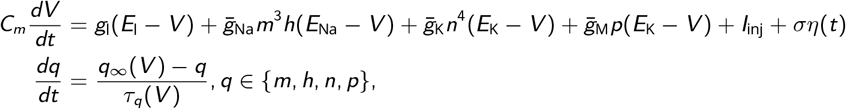

where *V* is the membrane potential, *C_m_* is the membrane capacitance, *g*_l_ is the leak conductance, *E*_l_ is the membrane reversal potential, 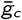 is the density of channels of type *c* (Na^+^, K^+^, M), *E_c_* is the reversal potential of *c*, (*m*, *h*, *n*, *p*) are the respective channel gating kinetic variables, and *ση*(*t*) is the intrinsic neural noise. The right hand side of the voltage dynamics is composed of a leak current, a voltage-dependent Na^+^ current, a delayed-rectifier K^+^ current, a slow voltage-dependent K^+^ current responsible for spike-frequency adaptation, and an injected current *I*_inj_. Channel gating variables *q* have dynamics fully characterized by the neuron membrane potential *V*, given the respective steady-state *q*_∞_(*V*) and time constant *τ_q_* (*V*) (details in [65]). Two additional parameters are implicit in the functions *q*_∞_(*V*) and *τ_q_*(*V*): *V_T_* adjusts the spike threshold through *m_∞_*, *h_∞_*, *n_∞_*, *τ_m_*, *τ_h_* and *τ_n_*; *τ*_max_ scales the time constant of adaptation through *τ_p_*(*V*) (details in [65]). We set *E*_Na_ = 53 mV and *E*_K_ = −107 mV, similar to the values used for simulations in Allen Cell Types Database (http://help.brain-map.org/download/attachments/8323525/BiophysModelPeri.pdf).

We applied SNPE to infer the posterior over 8 parameters (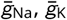, *g*_l_, 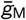, *τ*_max_, *V_T_*, *σ*, *E*_l_), given 7 voltage features (number of spikes, mean resting potential, standard deviation of the resting potential, and the first 4 voltage moments, mean, standard deviation, skewness and kurtosis).

The prior distribution over the parameters was uniform,

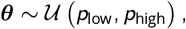

where *p*_low_ = [0.5, 10^−4^, 10^−4^, 10^−4^, 50, 40, 10^−4^, 35] and *p*_high_ = [80, 15, 0.6, 0.6, 3000, 90, 0.15, 100]. These ranges are similar to the ones obtained in [65].

For inference in simulated data, we used a single round of 100000 simulations (***θ*** ∈ ℝ^8^, **x** ∈ ℝ^11^). The density estimator was a masked autoregressive flow (MAF) [40] with five MADES with [50,50] hidden units each.

For the inference on in vitro recordings from mouse cortex (Allen Cell Types Database, https://celltypes.brain-map.org/data), we selected 8 recordings corresponding to spiny neurons with at least 10 spikes during the current-clamp stimulation. The respective cell identities and sweeps are: (518290966,57), (509881736,39), (566517779,46), (567399060,38), (569469018,44), (532571720,42), (555060623,34), (534524026,29). For each recording, SNPE-B was run for 2 rounds with 125000 Hodgkin–Huxley simulations each, and the posterior was approximated by a mixture of two Gaussians. In this case, the density estimator was composed of two fully connected layers of 100 units each.

### Comparison with genetic algorithm

We compared SNPE posterior with a state-of-the-art genetic algorithm (Indicator Based Evolutionary Algorithm IBEA, [70, 71] from the BluePyOpt package [31]), in the context of the Hodgkin-Huxley model with 8 parameters and 7 features (Supplementary Fig. 9). For each Hodgkin-Huxley model simulation *i* and summary feature *j*, we used the following objective score:

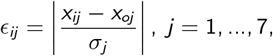

where *x_ij_* is the value of summary feature *j* for simulation *i*, *x_oj_* is the observed summary feature *j*, and *σ_j_* is the standard deviation of the summary feature *j* computed across 1000 previously simulated datasets. IBEA outputs the hall-of-fame, which corresponds to the 10 parameter sets with the lowest sum of objectives 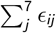. We ran IBEA with 100 generations and an offspring size of 1000 individuals, corresponding to a total of 100000 simulations.

### Circuit model of the crustacean stomatogastric ganglion

We used extracellular nerve recordings made from the stomatogastric motor neurons that principally comprise the triphasic pyloric rhythm in the crab *Cancer borealis* [74]. The preparations were decentralized, i.e. the axons of the descending modulatory inputs were severed. The data was recorded at a temperature of 11 °C. See [74] for full experimental details.

We simulated the circuit model of the crustacean stomatogastric ganglion by adapting a model described in [7]. The model is composed of three single-compartment neurons, AB/PD, LP, and PD, where the electrically coupled AB and PD neurons are modeled as a single neuron. Each of the model neurons contains 8 currents, a Na^+^ current *I*_Na_, a fast and a slow transient Ca^2+^ current *I*_CaT_ and *I*_CaS_, a transient K^+^ current *I*_A_, a Ca^2+^-dependent K^+^ current *I*_KCa_, a delayed rectifier K^+^ current *I*_Kd_, a hyperpolarization-activated inward current *I*_H_, and a leak current *I*_leak_. In addition, the model contains 7 synapses. As in [7], these synapses were simulated using a standard model of synaptic dynamics [138]. The synaptic input current into the neurons is given by *I*_s_ = *g*_s_*s*(*V*_post_ − *E*_s_), where *g*_s_ is the maximal synapse conductance, *V*_post_ the membrane potential of the postsynaptic neuron, and *E*_s_ the reversal potential of the synapse. The evolution of the activation variable *s* is given by

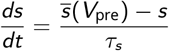

with

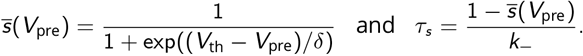

Here, *V*_pre_ is the membrane potential of the presynaptic neuron, *V*_th_ is the half-activation voltage of the synapse, *δ* sets the slope of the activation curve, and *k*_−_ is the rate constant for transmitter-receptor dissociation rate.

As in [7], two types of synapses were modeled since AB, LP, and PY are glutamatergic neurons whereas PD is cholinergic. We set *E_s_* = −70 mV and *k*_−_ = 1/40 ms for all glutamatergic synapses and *E_s_* = −80 mV and *k*_−_ = 1/100 ms for all cholinergic synapses. For both synapse types, we set *V*_th_ = −35 mV and *δ* = 5 mV.

For each set of membrane and synaptic conductances, we numerically simulated the rhythm for 10 seconds with a step size of 0.025 ms. To make the model stochastic, at each time step, we added Gaussian noise with a standard deviation of 0.001 mV to the input of each neuron.

We applied SNPE to infer the posterior over 24 membrane parameters and 7 synaptic parameters, i.e. 31 parameters in total. The 7 synaptic parameters were the maximal conductances *g*_s_ of all synapses in the circuit, each of which is varied uniformly in logarithmic domain from 0.01 nS to 1000 nS, with an exception of the synapse from AB to LP, which is varied uniformly in logarithmic domain from 0.01 nS to 10000 nS. The membrane parameters were the maximal membrane conductances for each of the neurons. The membrane conductances were varied over an extended range of previously reported values [7], which led us to the uniform prior bounds *p*_low_ = [0, 0, 0, 0, 0, 25, 0, 0] mS cm^−2^ and *p*_high_ = [500, 7.5, 8, 60, 15, 150, 0.2, 0.01] mS cm^−2^ for the maximal membrane conductances of the AB neuron, *p*_low_ = [0, 0, 2, 10, 0, 0, 0, 0.01] mS cm^−2^ and *p*_high_ = [200, 2.5, 12, 60, 10, 125, 0.06, 0.04] mS cm^−2^ for the maximal membrane conductances of the LP neuron, and *p*_low_ = [0, 0, 0, 30, 0, 50, 0, 0] mS cm−2 and *p*_high_ = [600, 12.5, 4, 60, 5, 150, 0.06, 0.04] mS cm^−2^ for the maximal membrane conductances of the PY neuron. The order of the membrane currents was: [Na, CaT, CaS, A, KCa, Kd, H, leak].

We used the 15 summary features proposed by [7], and extended them by 3 additional features. The features proposed by [7] are 15 salient features of the pyloric rhythm, namely: cycle period *T* (s), AB/PD burst duration 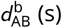, LP burst duration 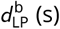, PY burst duration 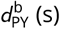, gap AB/PD end to LP start 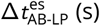, gap LP end to PY start 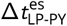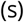, delay AB/PD start to LP start 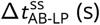, delay LP start to PY start 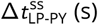, AB/PD duty cycle *d*_AB_, LP duty cycle *d*_LP_, PY duty cycle *d*_PY_, phase gap AB/PD end to LP start Δ*ϕ*_AB-LP_, phase gap LP end to PY start Δ*ϕ*_LP-PY_, LP start phase *ϕ*_LP_, and PY start phase *ϕ*_PY_. Note that several of these values are only defined if each neuron produces rhythmic bursting behavior. In addition, for each of the three neurons, we used one feature that describes the maximal duration of its voltage being above −30 mV. We did this as we observed plateaus at around −10 mV during the onset of bursts, and wanted to distinguish such activity traces from others. If the maximal duration was below 5 ms, we set this feature to 5 ms. To extract the summary features from the observed experimental data, we first found spikes by searching for local maxima above a hand-picked voltage threshold, and then extracted the 15 above described features. We set the additional 3 features to 5 ms.

We used SNPE to infer the posterior distribution over the 18 summary features from experimental data. For inference, we used a single round with 18.5 million samples, out of which 174,000 samples contain bursts in all neurons. We therefore used these 174,000 samples with well defined summary features for training the inference network (***θ*** ∈ ℝ^31^, **x** ∈ ℝ^18^). The density estimator was a masked autoregressive flow (MAF) [40] with five MADES with [200,400] hidden units each. The synaptic conductances were transformed into logarithmic space before training and for the entire analysis.

Previous approaches for fitting the STG circuit [7] first fit individual neuron features and reduce the number of possible neuron models [25], and then fit the whole circuit model. While powerful, this approach both requires the availability of single-neuron data, and cannot give access to potential compensation mechanisms between single-neuron and synaptic parameters. Unlike [7], we apply SNPE to directly identify the full 31 dimensional parameter space without requiring experimental measurements of each individual neuron in the circuit. Despite the high-dimensional parameter space, SNPE can identify the posterior distribution using 18 million samples, whereas a direct application of a full-grid method would require 4.65 · 10^21^ samples to fill the 31 dimensional parameter space on a grid with five values per dimension.

### Finding paths in the posterior

In order to find directions of robust network output, we searched for a path of high posterior probability. First, as in [7], we aimed to find 2 similar model outputs with disparate parameters. To do so, we sampled from the posterior and searched for 2 parameter sets whose summary features were within 0.1 standard deviations of all 174,000 samples from the observed experimental data, but that had strongly disparate parameters from each other. In the following, we denote the obtained parameter sets by ***θ***_*s*_ and ***θ***_*g*_.

Second, in order to identify whether network output can be maintained along a continuous path between these 2 samples, we searched for a connection in parameter space lying in regions of high posterior probability. To do so, we considered the connection between the samples as a path and minimize the following path integral:

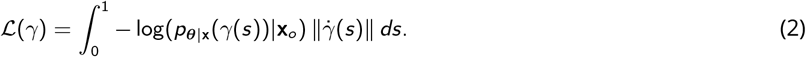

To minimize this term, we parameterized the path *γ*(*s*) using sinusoidal basis-functions with coefficients *α*_*n,k*_ :

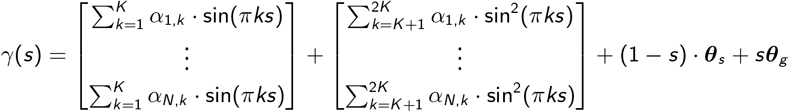

These basis functions are defined such that, for any coefficients *α*_*n,k*_, the starting and end points of the path are exactly the two parameter sets defined above:

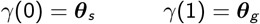

With this formulation, we have framed the problem of finding the path as an unconstrained optimization problem over the parameters *α*_*n,k*_. We can therefore minimize the path integral *L* using gradient descent over *α*_*n,k*_. For numerical simulations, we approximated the integral in equation 2 as a sum over 80 points along the path and use 2 basis functions for each of the 31 dimensions, i.e. *K* = 2.

In order to demonstrate the sensitivity of the pyloric network, we aimed to find a path along which the circuit output quickly breaks down. For this, we picked a starting point along the high-probability path and then minimize the posterior probability. In addition, we enforced that the orthogonal path lies within an orthogonal disk to the high-probability path, leading to the following constrained optimization problem:

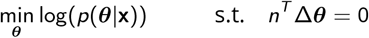

where *n* is the tangent vector along the path of high probability. This optimization problem can be solved using the gradient projection method [139]:

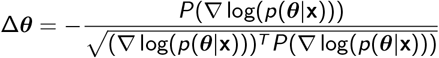

with projection matrix 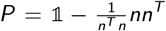 and 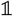 indicating the identity matrix. Each gradient update is a step along the orthogonal path. We let the optimization run until the distance along the path is 1/27 of the distance along the high-probability path.

### Identifying conditional correlations

In order to investigate compensation mechanisms in the STG, we compared marginal and conditional correlations. For the marginal correlation matrix in Fig. 6b, we calculated the Pearson correlation coefficient based on 1.26 million samples from the posterior distribution *p*(***θ***|**x**). To find the 2-dimensional conditional distribution for any pair of parameters, we fixed all other parameters to values taken from an arbitrary posterior sample, and varied the remaining 2 on an evenly spaced grid with 50 points along each dimension, covering the entire prior space. We evaluated the posterior distribution at every value on this grid. We then calculated the conditional correlation as the Pearson correlation coefficient over this distribution. For the 1-dimensional conditional distribution, we varied only 1 parameter and kept all others fixed. Lastly, in Fig. 6d, we sampled 500 parameter sets from the posterior, computed the respective conditional posteriors and conditional correlation matrices, and took the average over the conditional correlation matrices.

## Supplementary material

### Supplementary figures

**Supplementary Figure 1.**
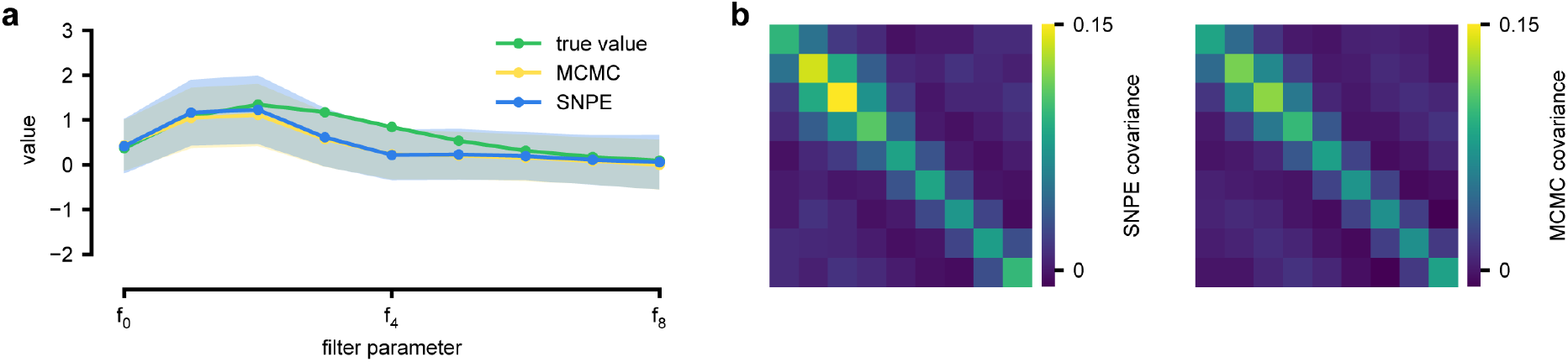
Comparison between SNPE-estimated posterior and reference posterior (obtained via MCMC) on LN model. (a) Posterior mean ± one standard deviation of temporal filter (receptive field) from SNPE posterior (SNPE, blue) and reference posterior (MCMC, yellow). (b) Full covariance matrices from SNPE posterior (left) and reference (MCMC, right).

**Supplementary Figure 2.**
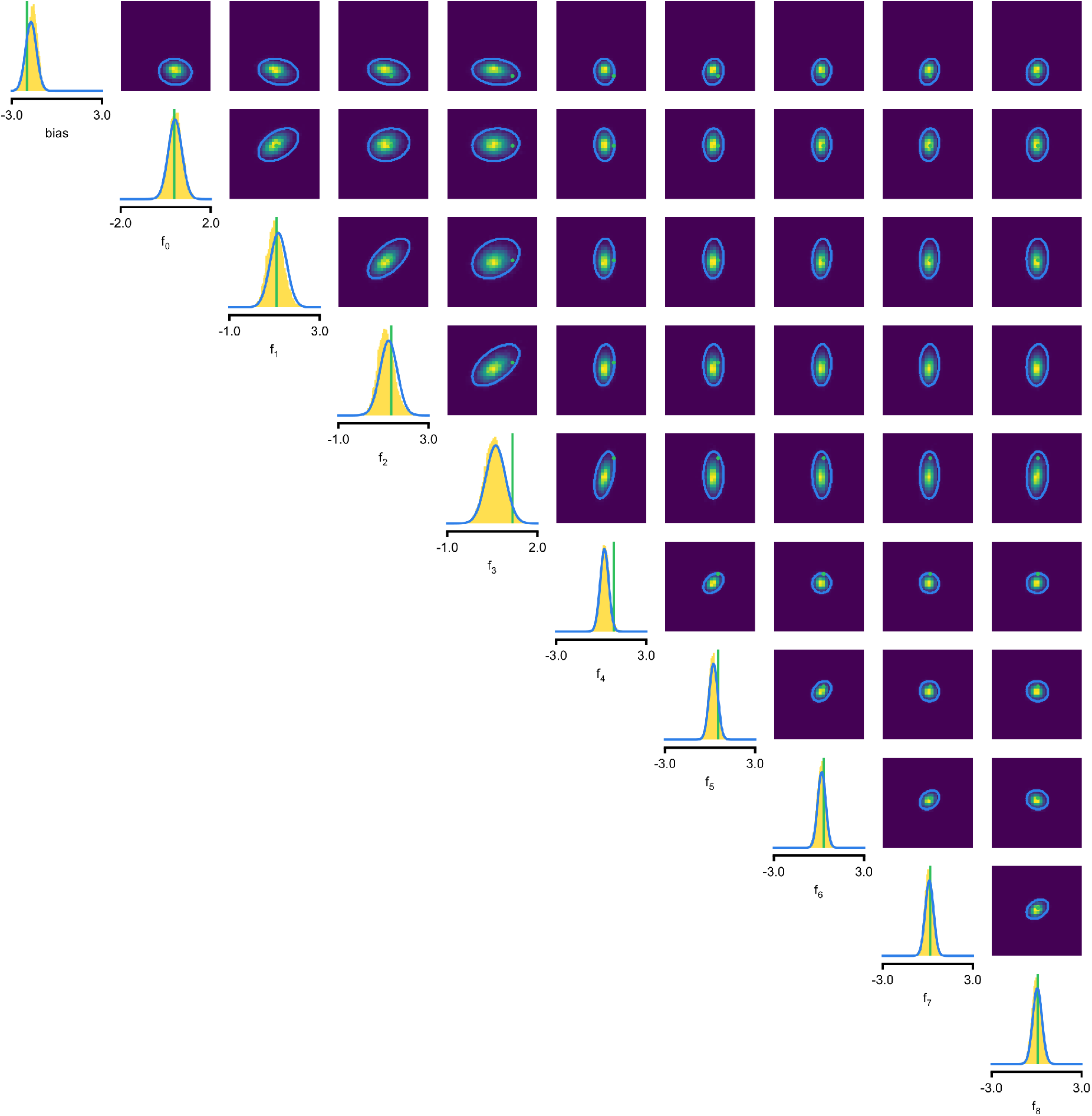
Full posterior for LN model. In green, ground-truth parameters. Marginals (blue lines) and 2D marginals for SNPE (contour lines correspond to 95% of the mass) and MCMC (yellow histograms).

**Supplementary Figure 3.**
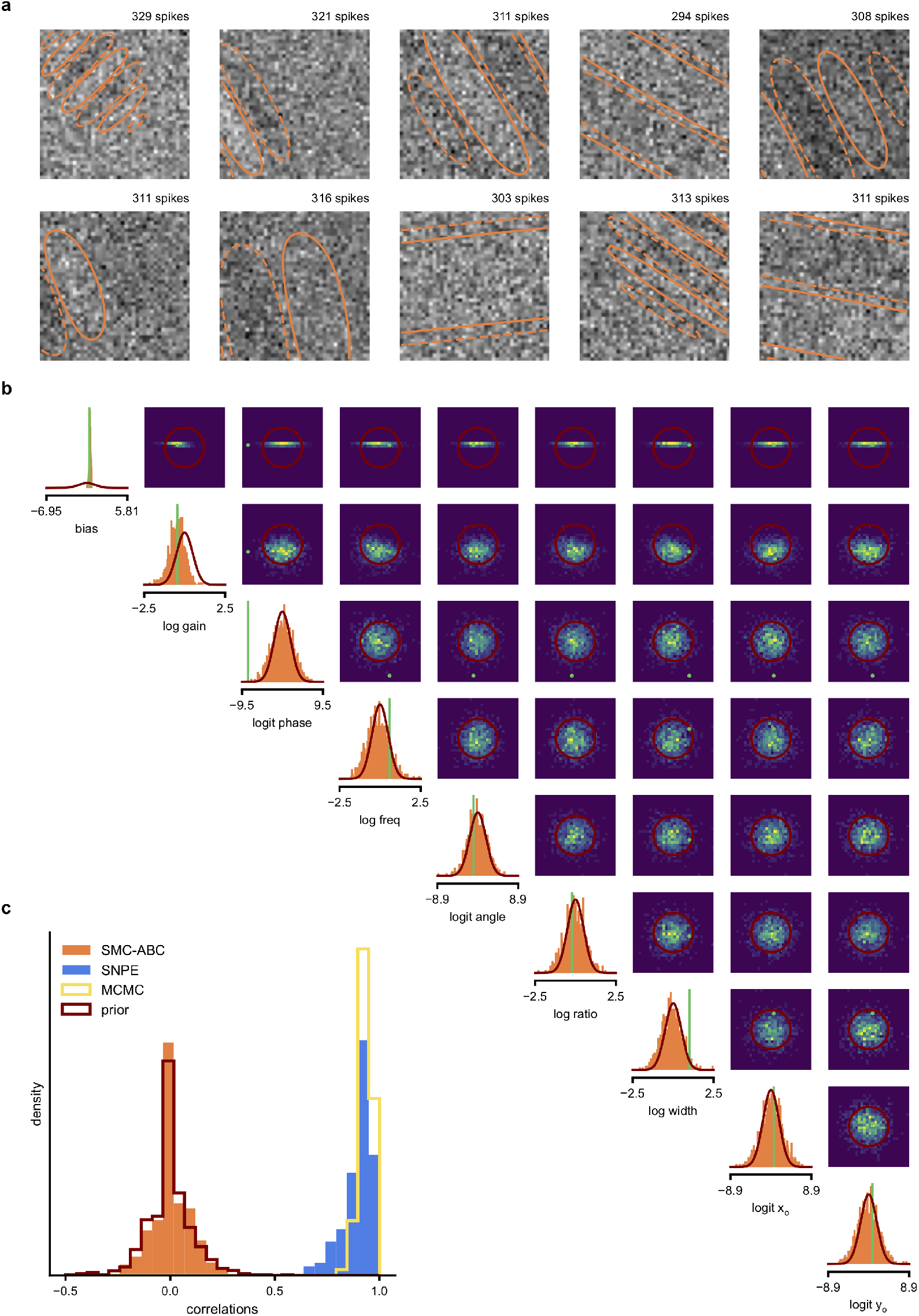
SMC-ABC posterior estimate for Gabor GLM receptive field model. (a) Spike-triggered averages (STAs) and spike counts with closest distance *d* (*x_o_*, *x_i_*) to the observed data *x_o_* out of 10000 simulations with *θ_i_* sampled from the prior. Spike counts are comparable to the observed data (*x_o_* : 299 spikes), but receptive fields (contours) are not well constrained. (b) Results for SMC-ABC with 10^6^ simulations total. Histograms of 1000 particles (orange) returned in the final iteration of SMC-ABC, compared to prior (red contour lines) and ground-truth parameters (green). Distributions over (log-/logit-)transformed parameters, axis limits scaled to mean ± 3 standard deviations of the prior. (c) Correlations between ground-truth receptive field and receptive fields sampled from SMC-ABC posterior (orange), SNPE posterior (blue), reference MCMC posterior (yellow) and prior (red). The SNPE-estimated receptive fields are almost as good as those of the reference posterior, the SMC-ABC estimated ones no better than the prior.

**Supplementary Figure 4.**
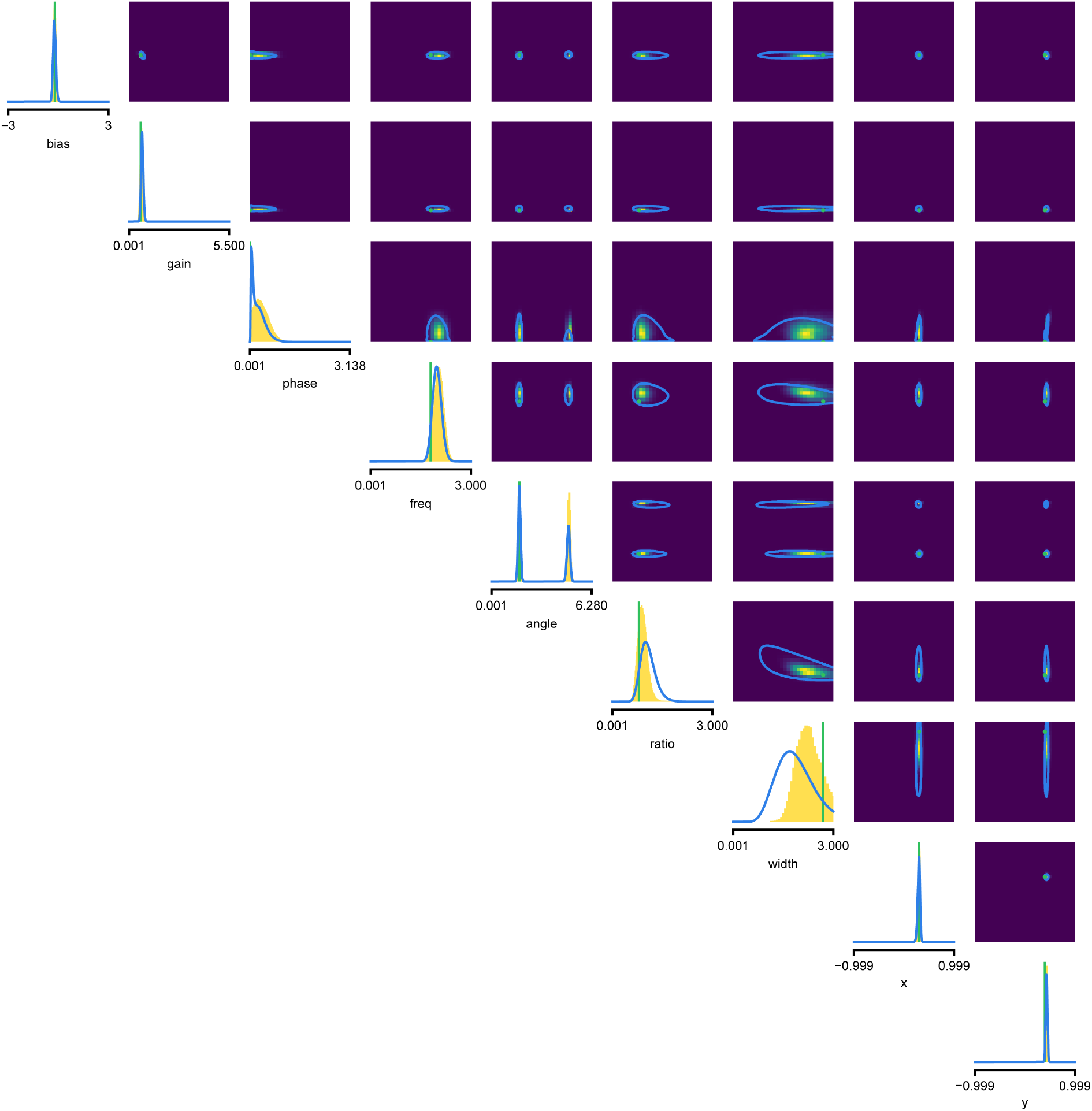
Full posterior for Gabor GLM receptive field model. SNPE posterior estimate (blue lines) compared to reference posterior (MCMC, histograms). Ground-truth parameters used to simulate the data in green. We depict the distributions over the original receptive field parameters, whereas we estimate the posterior as a Gaussian mixture over transformed parameters, see Methods for details. We find that a (back-transformed) Gaussian mixture with four components approximates the posterior well in this case.

**Supplementary Figure 5.**
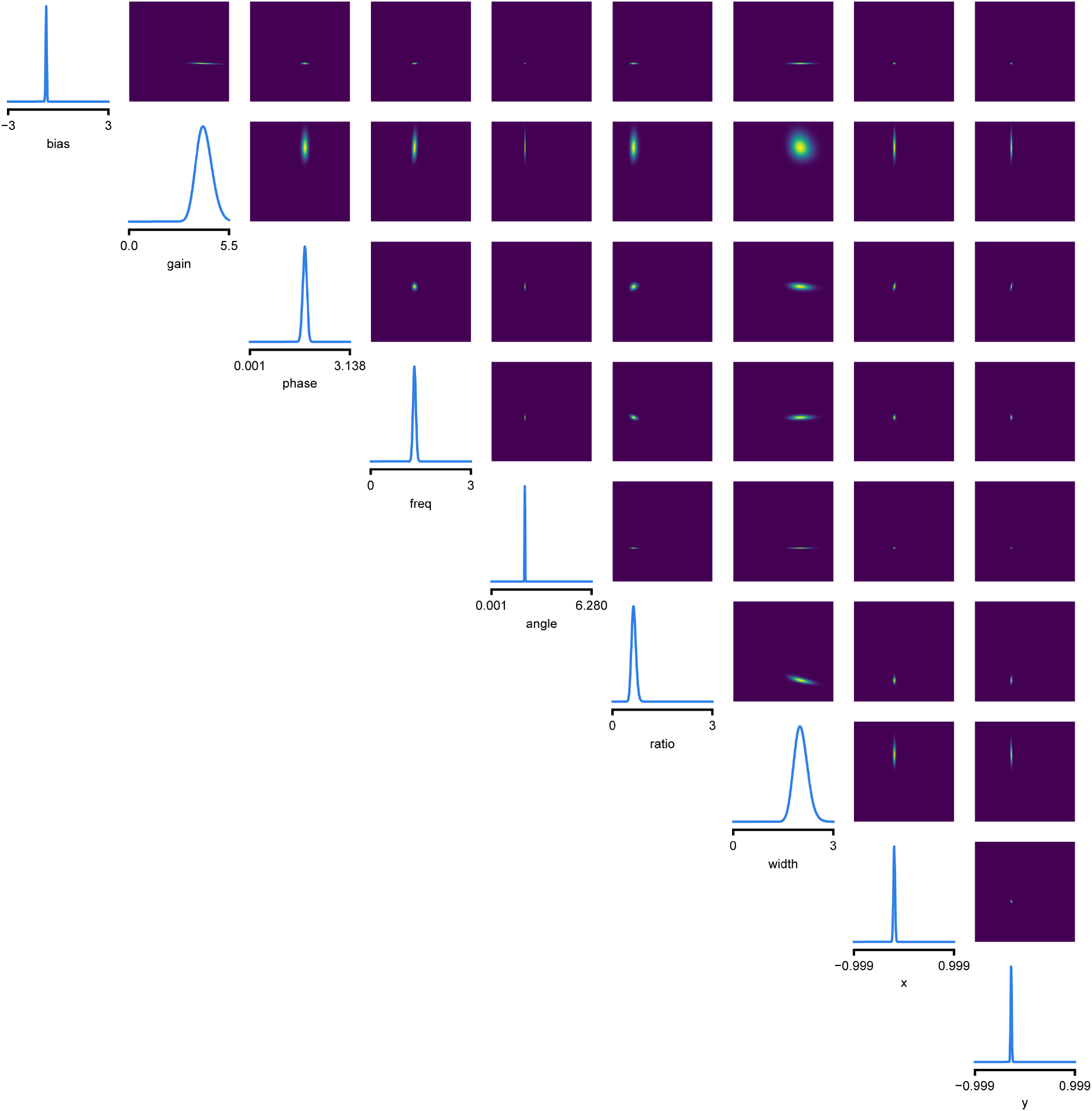
Full posterior for Gabor LN receptive field model on V1 recordings. We depict the distributions over the receptive field parameters, derived from the Gaussian mixture over transformed-parameters (see Methods for details).

**Supplementary Figure 6.**
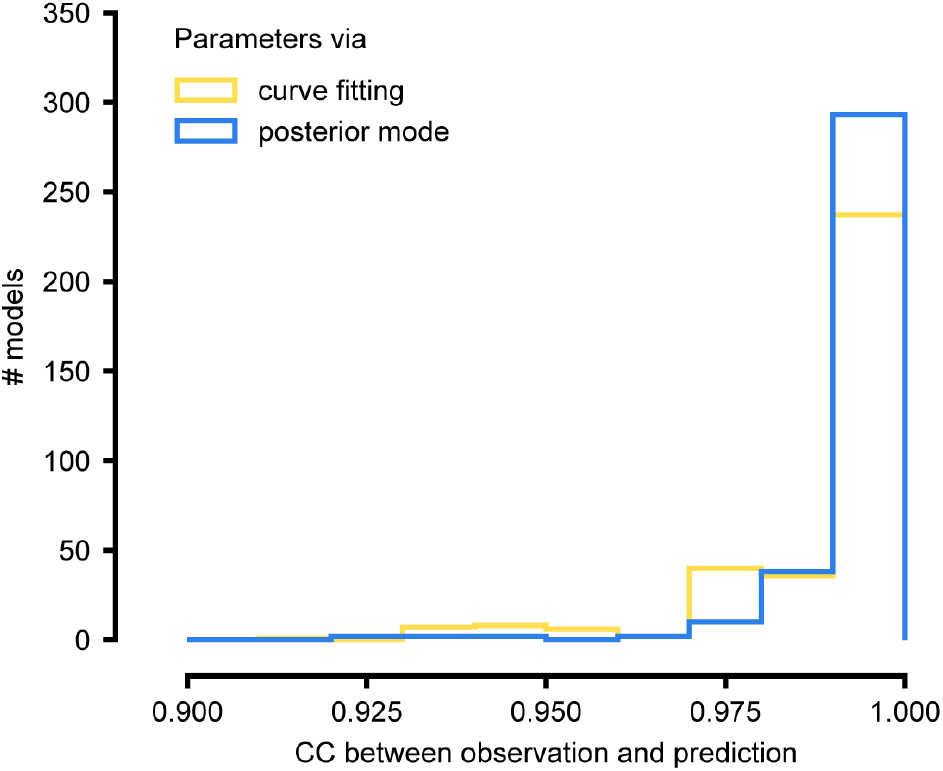
Summary results on ICG channel models, and comparison with direct fits. We generate predictions either with the posterior mode (blue) or with parameters obtained by directly fitting steady-state activation and time-constant curves (yellow). We calculate the correlation coefficient (CC) between observation and prediction. The distribution of CCs is similar for both approaches.

**Supplementary Figure 7.**
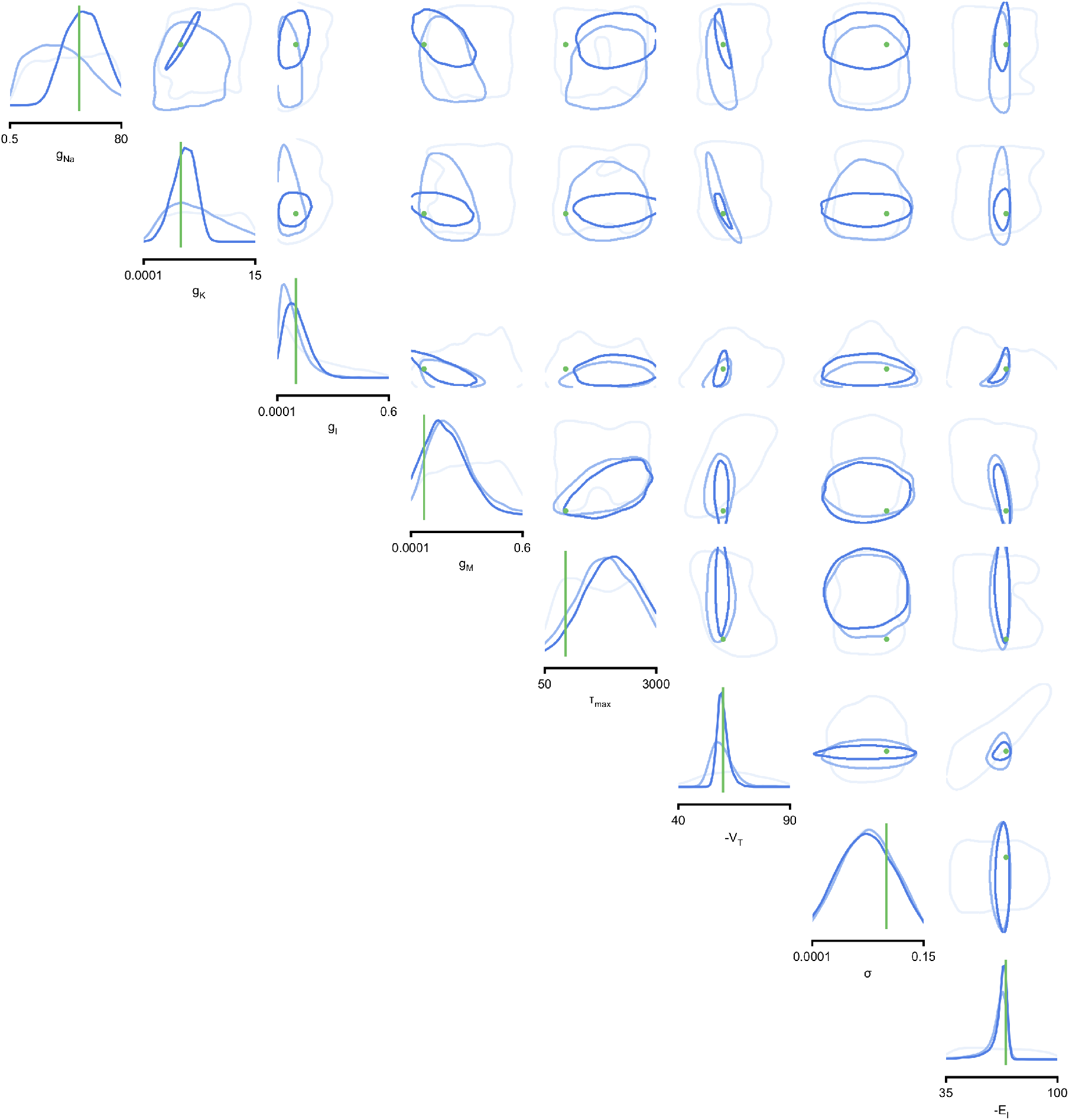
Full posteriors for Hodgkin-Huxley model for 1, 4 and 7 features. Images show the pairwise marginals for 7 features. Each contour line corresponds to 68% density mass for a different inferred posterior. Light blue corresponds to 1 feature and dark blue to 7 features.

**Supplementary Figure 8.**
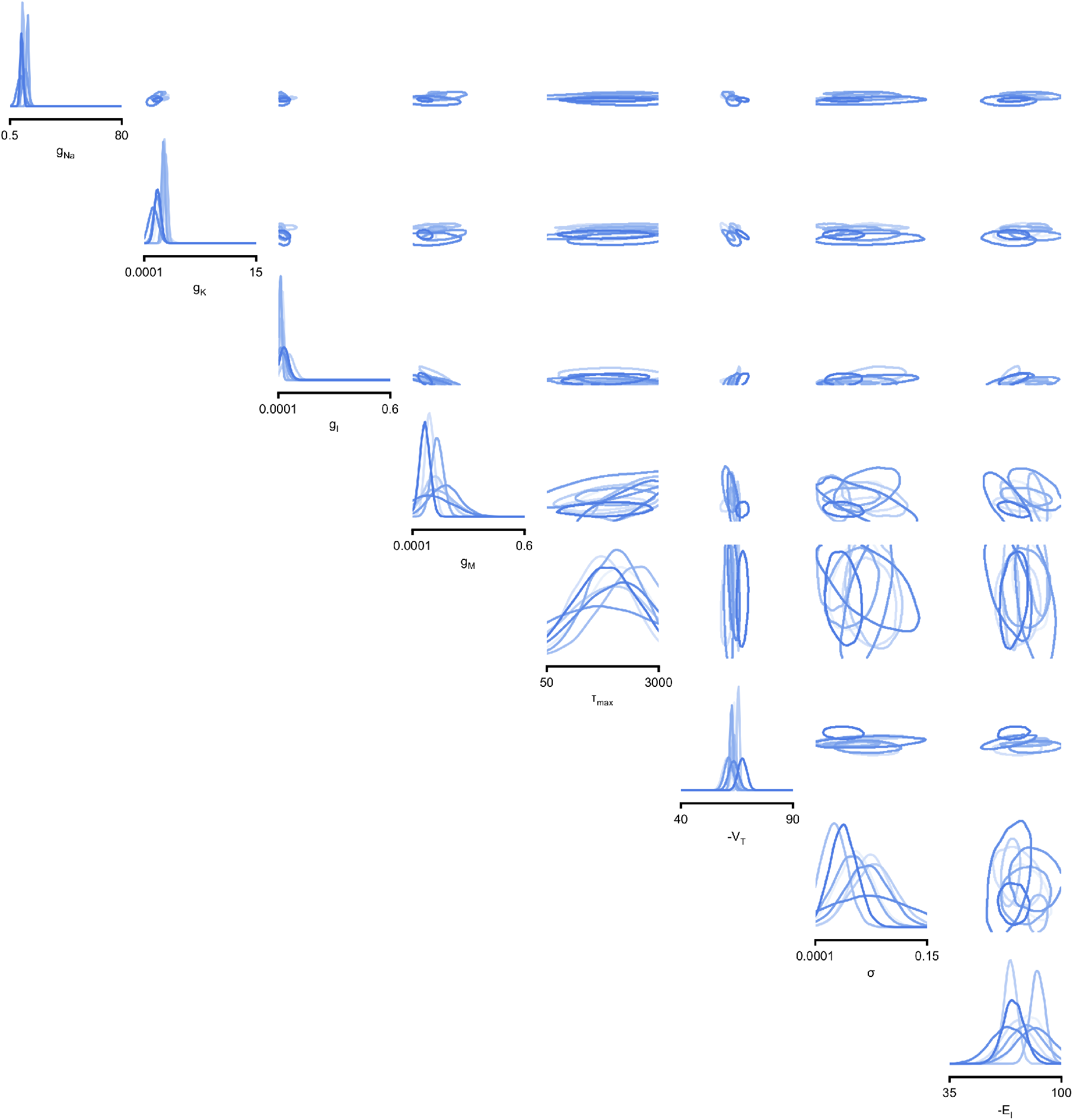
Full posteriors for Hodgkin-Huxley model on 8 different recordings from Allen Cell Type Database. Images show the pairwise marginals for 7 features. Each contour line corresponds to 68% density mass for a different inferred posterior.

**Supplementary Figure 9.**
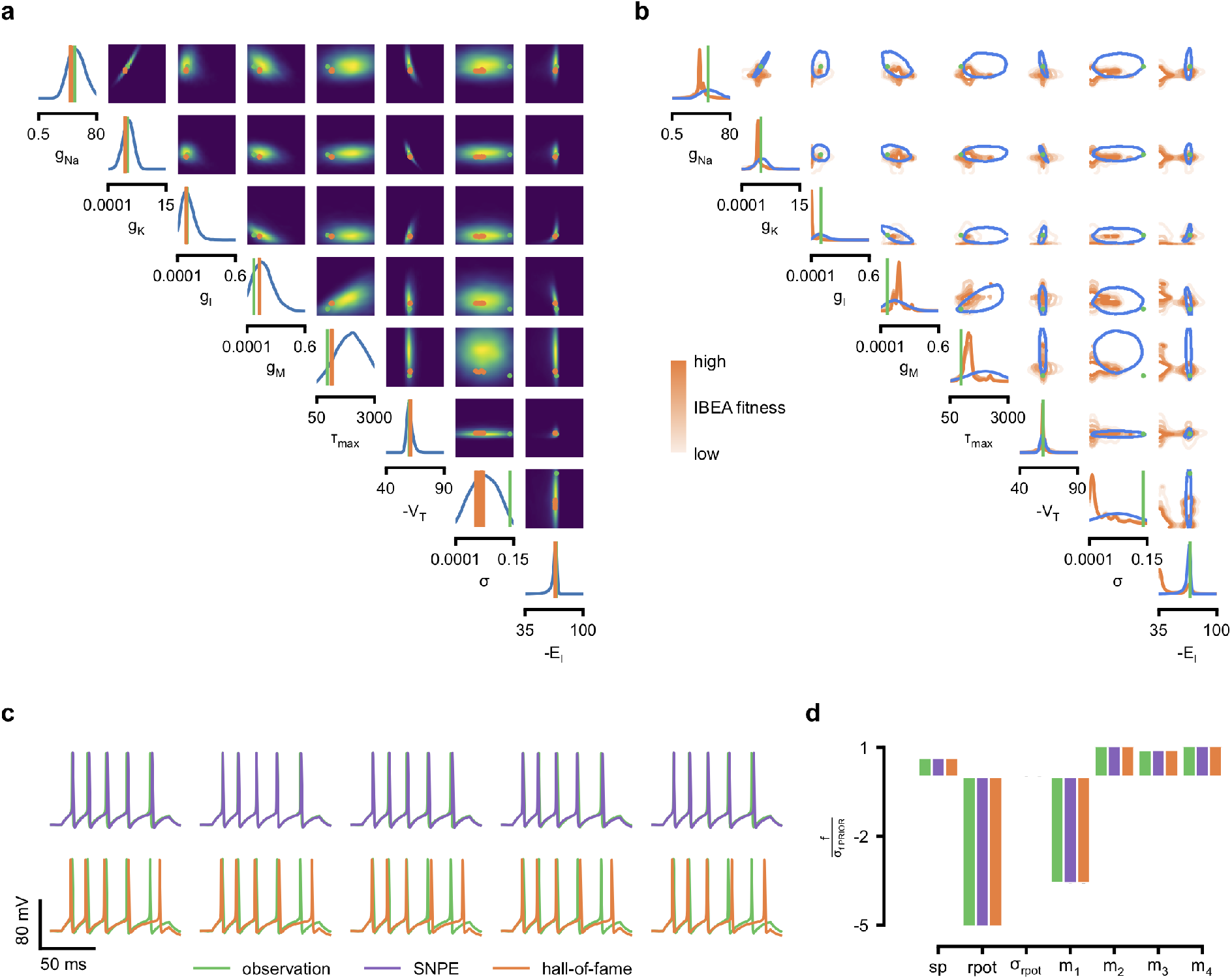
Comparison between SNPE posterior and IBEA samples for Hodgkin-Huxley model with 8 parameters and 7 features. (a) Full SNPE posterior distribution. Ground truth parameters in green and IBEA 10 parameters with highest fitness (‘hall-of-fame’) in orange. (b) Blue contour line corresponds to 68% density mass for SNPE posterior. Light orange corresponds to IBEA sampled parameters with lowest IBEA fitness and dark orange to IBEA sampled parameters with highest IBEA fitness. This plot shows that, in general, SNPE and IBEA can return very different answers– this is not surprising, as both algorithms have different objectives, but this highlights that genetic algorithms do not in general perform statistical inference. (c) Traces for samples with high probability under SNPE posterior (purple), and for samples with high fitness under IBEA objective (hall-of-fame; orange traces). (d) Features for the desired output (observation), the mode of the inferred posterior (purple) and the best sample under IBEA objective (orange). Each voltage feature is normalized by *σ*_f PRIOR_, the standard deviation of the respective feature of simulations sampled from the prior.

**Supplementary Figure 10.**
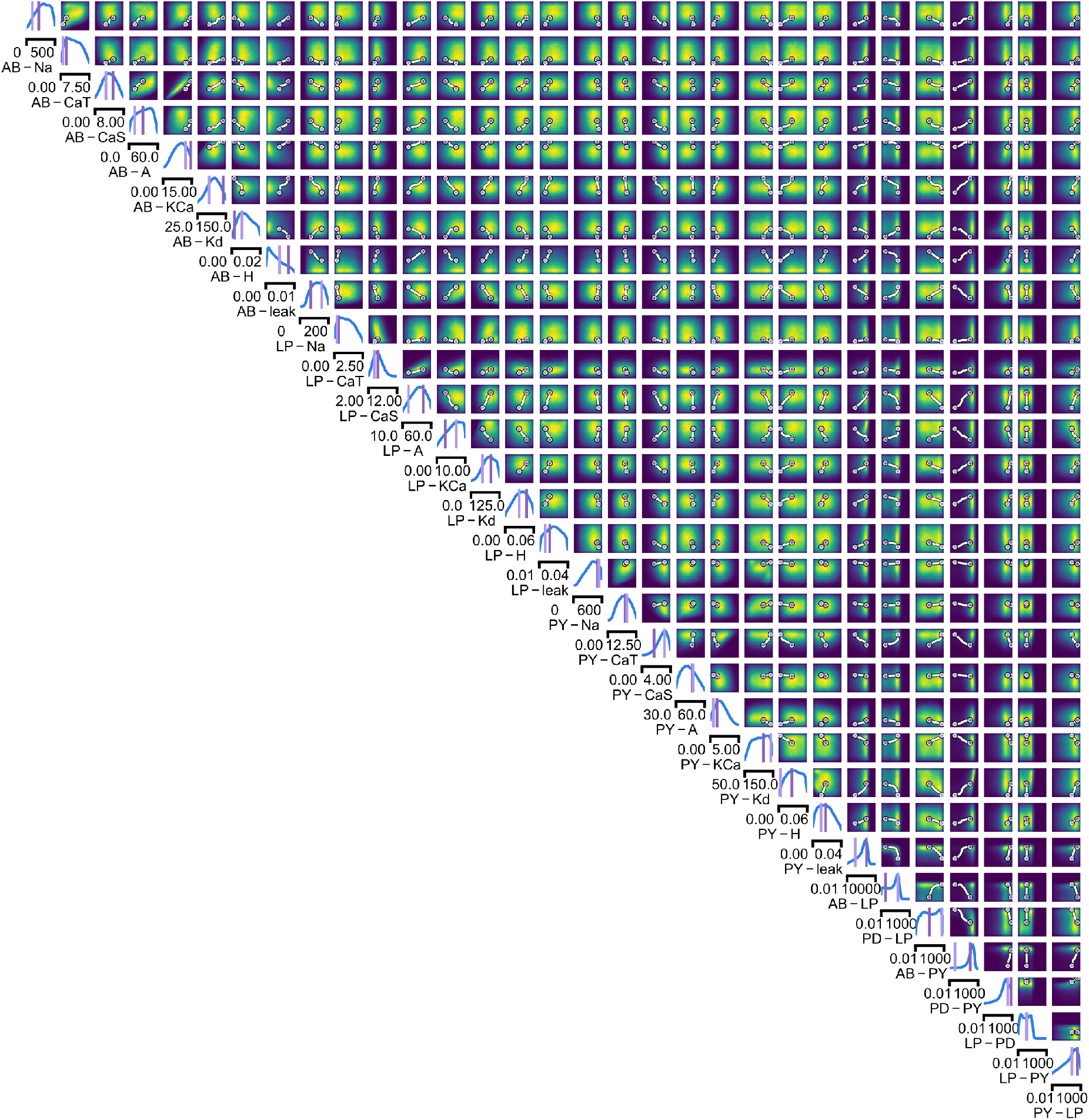
Full posterior for the stomatogastric ganglion over 24 membrane and 7 synaptic conductances. The first 24 dimensions depict membrane conductances (top left), the last 7 depict synaptic conductances (bottom right). All synaptic conductances are logarithmically spaced. Between two samples from the posterior with high posterior probability (purple dots), there is a path of high posterior probability (white).

**Supplementary Figure 11.**
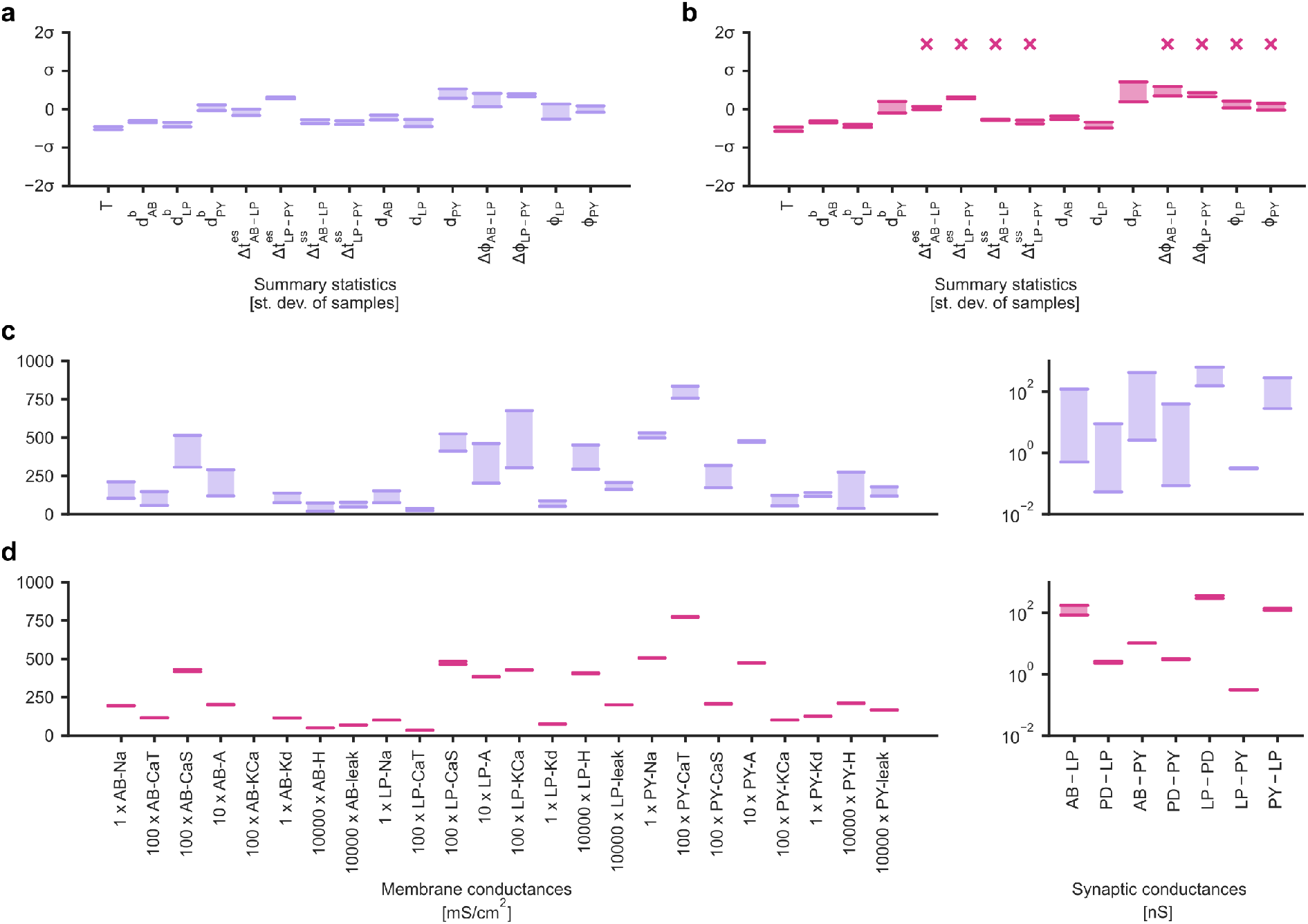
Identifying directions of sloppiness and stiffness in the pyloric network of the crustacean stomatogastric ganglion. (a) Minimal and maximal values of all summary statistics along the path lying in regions of high posterior probability, sampled at 20 evenly spaced points. Summary statistics change only little. The summary statistics are scaled with the standard deviation of the 170,000 bursting samples in the created dataset. (b) Summary statistics sampled at 20 evenly spaced points along the orthogonal path. The summary statistics show stronger changes than in panel a and, in particular, often could not be defined because neurons bursted irregularly, as indicated by an ‘x’ above barplots. (c) Minimal and maximal values of the circuit parameters along the path lying in regions of high posterior probability. Both membrane conductances (left) and synaptic conductances (right) vary over large ranges. Axes as in panel (d). (d) Circuit parameters along the orthogonal path. The difference between the minimal and maximal value is much smaller than in panel (c).

**Supplementary Figure 12.**
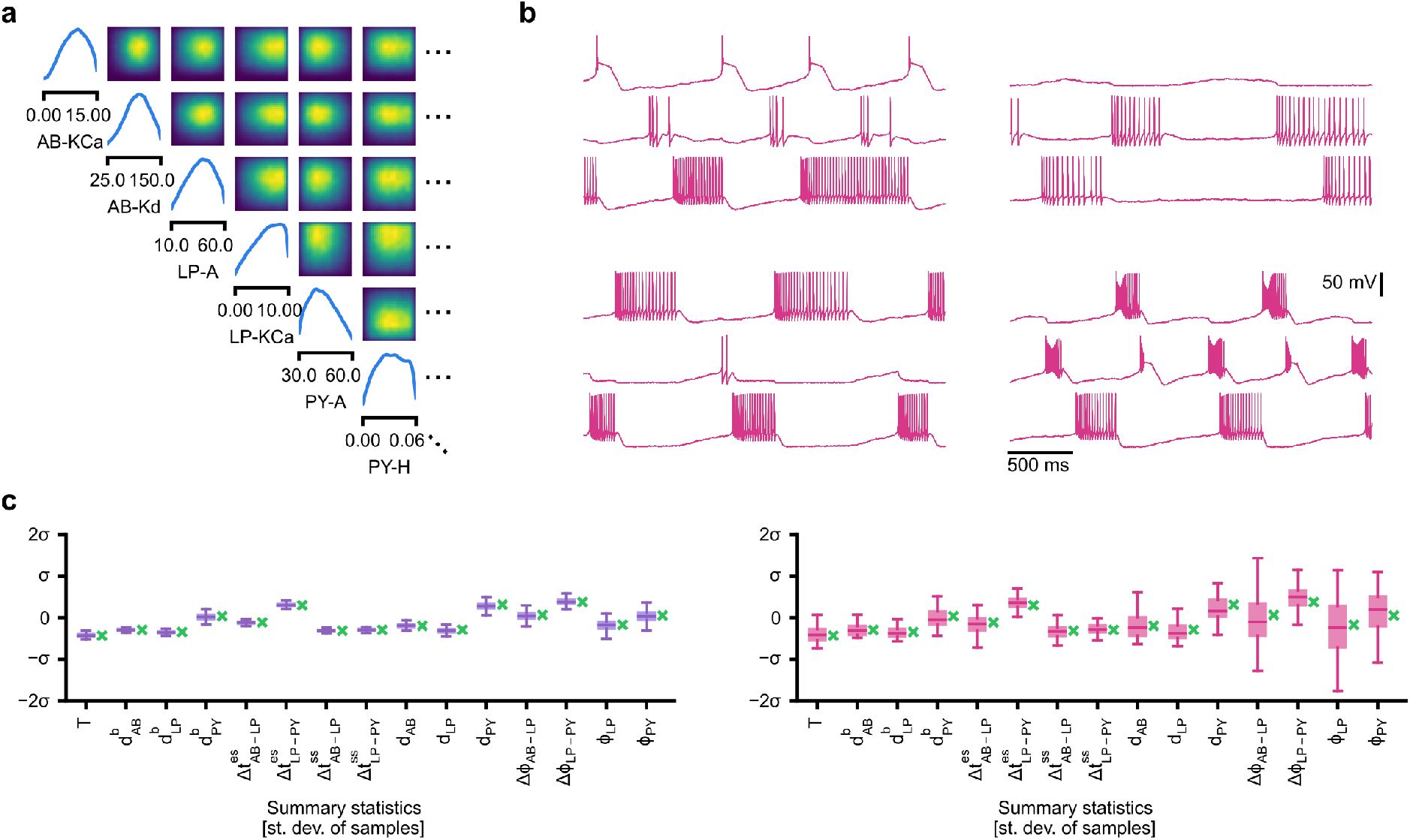
Evaluating circuit configurations in which parameters have been sampled independently. (a) Factorized posterior, i.e. posterior obtained by sampling each parameter independently from the associated marginals. Many of the pairwise marginals look similar to the full posterior shown in Supplementary Fig. 10, as the posterior correlations are low. (b) Samples from the factorized posterior– only a minority of these samples produce pyloric activity, highlighting the significance of the posterior correlations between parameters. (c) Left: summary features for 500 samples from the posterior. Boxplot for samples where all summary features are well-defined (80 % of all samples). Right: summary features for 500 samples from the factorized posterior. Only 23 % of these samples have well-defined summary features. The summary features from the factorized posterior have higher variation than the posterior ones. Summary features are normalized using the mean and standard deviation of all samples in our training dataset obtained from prior samples. The boxplots indicate the maximum, 75% quantile, median, 25% quantile, and minimum. The green ‘x’ indicates the value of the experimental data (the observation, shown in figure 5B).

